# Proximity directs microbial transglutaminase site selectivity in native antibody modification

**DOI:** 10.64898/2026.08.18.745451

**Authors:** Riko Nishioka, Koki Murozono, Yoshirou Kawaguchi, Michio Kimura, Shun Sakuraba, Noritaka Hashii, Akinobu Senoo, Jose M.M. Caaveiro, Mitsuo Umetsu, Noriho Kamiya

## Abstract

Site-specific protein modification allows diverse functionalities to be introduced while minimizing perturbations to the protein structure and activity. Considerable efforts have been made to achieve site-specific modification of native proteins to overcome the heterogeneity resulting from conventional stochastic Lys or Cys modification. We have previously achieved the selective modification of Lys65 in a native immunoglobulin G1 (IgG1) antibody (trastuzumab) using EzMTG-pG(Fab), which is an engineered zymogen of microbial transglutaminase (EzMTG) fused to a Fab-binding protein G [pG(Fab)]. However, this approach cannot be widely applied to different types of IgG antibodies. Here, we designed pG(Fab)-EzMTG by fusing pG(Fab) to the *N*-terminus of EzMTG. Notably, switching the fusion partners dramatically altered the IgG modification site from Lys65 to Lys225, which is located in the hinge site of native IgG1 antibodies. This Lys225-selective labeling was applicable to different IgG1 antibodies. As a functional application, the cytotoxic drug monomethyl auristatin E (MMAE) was conjugated to Lys225 of trastuzumab, and the resulting antibody–drug conjugate exhibited antigen-specific cytotoxicity. These findings demonstrate that fusion-protein architecture determines site selectivity in proximity-directed enzymatic modification, providing a strategy for the site-specific functionalization of native antibodies.

## Introduction

The conjugation of fluorescent probes, drugs, and other functional molecules to proteins expands their functionality and enables diverse applications in biological research, diagnostics, and therapeutics. However, conventional bioconjugation methods generally offer limited control over the site and degree of modification. This results in heterogeneous mixtures of protein conjugates with variable biological activities.^1, 2, 3^ Accordingly, site-specific protein modification provides a powerful means of introducing functionalities into proteins while preserving their native structures and activities.^4^

Antibodies are important but challenging targets for site-specific protein modification. A typical IgG has approximately 40 solvent-accessible Lys residues, and modification based on their intrinsic reactivity results in heterogeneous populations with different conjugation sites and degrees of modification.^5^ The conjugation site can considerably influence the stability, efficacy, pharmacokinetics, and safety profiles. This is particularly important in antibody–drug conjugates (ADCs), which combine the high target specificity of monoclonal antibodies with the potent cytotoxic activity of small-molecule payloads.^6^ However, most ADCs have been generated by exploiting the reactivity of Lys or Cys residues on the antibody surface, resulting in heterogeneous ADC populations.^7^ This heterogeneity can lead to reduced efficacy because of the rapid clearance of species with a high drug-to-antibody ratio (DAR), as well as compromising reproducibility and manufacturability. Consequently, considerable efforts have been directed towards developing site-specific antibody modification strategies.^8, 9, 10^

Common strategies for site-specific antibody modification include the introduction of additional Cys residues, as exemplified by THIOMAB technology^11^, or non-canonical amino acids into antibodies.^12, 13, 14^ Enzyme-mediated strategies, such as those employing sortase A and transglutaminases, have also proven effective for site-specific antibody modification.^15, 16, 17^ Sortase A recognizes a *C*-terminal Leu-Pro-X-Thr-Gly (LPXTG) motif and cleaves the Thr–Gly bond to ligate oligoglycine-containing substrates (e.g., GGG)^18^, enabling site-specific conjugation when a GGG sequence is introduced at the antibody *C*-terminus.^19^ Transglutaminases catalyze acyl transfer reactions between the γ-carboxamide group of the side chain of Gln residues and the ε-amino group of Lys side chains, as well as acyl transfer reactions between Gln residues and primary amines within proteins.^20^ The use of microbial transglutaminase (MTG) is particularly attractive because MTG operates under mild conditions without requiring cofactors^21^, and this transglutaminase has therefore been widely used in the food industry and in ADC production for site-specific antibody modification. Several studies have reported the site-specific modification of antibodies following deglycosylation^22, 23^ or genetic mutagenesis.^24, 25, 26^ However, these approaches require antibody pretreatment or engineering, which may affect antibody expression yields and compromise native antibody functions.^27^ Consequently, there has been increasing attention directed toward developing tag-free enzymatic strategies for the site-specific modification of native antibodies.^16^

Tag-free antibody modification using MTG has been reported. Dickgiesser et al. have demonstrated that an MTG variant, in which a Gly residue near the active-site loop was substituted with Ser, enabled selective modification of the native IgG residue Gln295.^28^ Similarly, Wehrmüller et al. have achieved site-selective modification of Gln295 on native IgG by performing MTG-mediated conjugation in the presence of short cationic peptides that enhanced local reactivity.^29^ In addition to MTG-based approaches, tag-free modification of native antibodies has been reported using other enzymes, such as horseradish peroxidase^30^ and lipoic acid ligase A.^31, 32^

Recently, alternative strategies based on the proximity effect have been used to enable site-specific labeling of native antibodies.^33, 34^ The proximity effect refers to the recruitment of crosslinking reagents or enzymes to a target amino acid residue through ligands that bind to defined sites on antibodies, such as Fc-affinity peptides or antibody-binding proteins. By localizing reactive reagents or catalysts near the antibody, these ligand-guided proximity approaches achieve site-selective modification without requiring genetic engineering. Representative examples include pClick^35^ and chemical conjugation by affinity peptide (CCAP)^36^ in which Fc-affinity peptides bearing Lys-reactive groups form covalent bonds with nearby Lys residues, enabling selective modification of specific sites within native antibodies. More recently, ‘traceless’ affinity-based approaches have been developed to minimize structural changes in antibodies.^5^ The AJICAP method employs an Fc-III peptide linked via a disulfide-containing linker to introduce free thiols into native antibodies to produce ADCs with uniform DAR values. As the peptide used for proximity targeting is ultimately removed, AJICAP is a traceless ADC preparation strategy.^37, 38^ Proximity-based methods using antibody-binding proteins as ligands have also been demonstrated by fusing reactive probes^39^ or sortase A variants^40^ to antibody-binding proteins, enabling site-specific antibody modification. As the antibody-binding protein fusion enzymes can be removed from the final products, such approaches also provide ‘traceless’ conjugation strategies.

Previously, we designed EzMTG-pG(Fab), a fusion construct comprising a protein G variant [pG(Fab)] that specifically binds to a Fab region fused to the *C*-terminus of an engineered zymogen of microbial transglutaminase (EzMTG^41^).^42^ This construct can be expressed as a soluble protein in *Escherichia coli*. We successfully achieved the site-specific modification of Lys65 of a native IgG (trastuzumab) with a fluorescent substrate using EzMTG-pG(Fab). Because pG(Fab) binds to the Fab domain, Lys65-selective modification was achieved for the Fab fragment of trastuzumab.^43^ However, as Lys65 is located close to the antigen-binding region, this site is not fully compatible with antibody species other than trastuzumab.

In the present study, we explored the selective modification of alternative Lys residues. Looking at the structural characteristics of pG(Fab), we found that the *N*- and *C*-termini exhibited distinct orientations, leading to the idea of relocating EzMTG from the *N*-terminus to the *C*-terminus of pG(Fab). EzMTG retains an *N*-terminal propeptide that facilitates the proper folding of MTG. This propeptide enables the introduction of functional domains at the *N*-terminus of MTG, a modification that has previously proven challenging. Thus, we designed a new construct, pG(Fab)-EzMTG, in which pG(Fab) was fused to the *N*-terminal side of EzMTG (**Fig. 1, Table S1**). Simply switching the orientation of the fusion construct redirected the predominant modification site from Lys65 to Lys225 in the hinge region of native IgG1 antibodies, without altering the catalytic domain of EzMTG. Lys225-selective modification was also applicable to multiple IgG1 antibodies. These results demonstrate that the spatial arrangement of the binding and catalytic domains is a key determinant of site selectivity in proximity-directed enzymatic modification, providing a strategy for the site-specific functionalization of native antibodies.

**Fig. 1.**
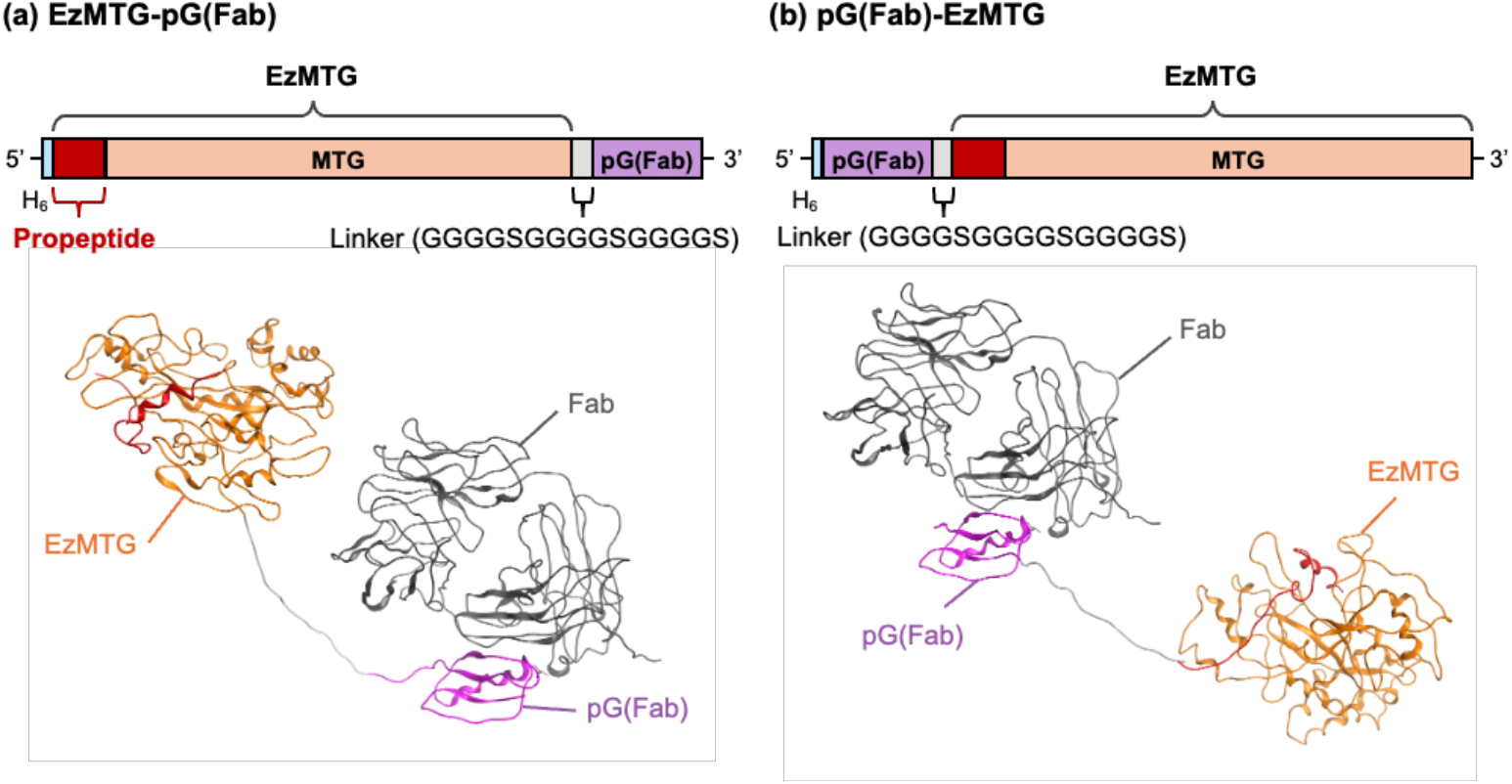
The pG(Fab)-EzMTG construct designed in the present study. Schematic representation of (a) EzMTG-pG(Fab) and (b) pG(Fab)-EzMTG. The three-dimensional structure of EzMTG-pG(Fab) and pG(Fab)-EzMTG. Trastuzumab (Fab), EzMTG, and pG(Fab) are shown in gray, orange, and purple, respectively. The PDB entries for the Fab-pG(Fab) complex and EzMTG are 1IGC.pdb and 3IU0.pdb, respectively.

## Results and Discussion

### Evaluation of the crosslinking activity of pG(Fab)-EzMTG

To evaluate the crosslinking activity of pG(Fab)-EzMTG, an IgG antibody labeling reaction was performed using a TAMRA-labeled Gln substrate (TAMRA-YPLQMRG-NH_2_, TAMRA-Q, Figure S1). As with EzMTG-pG(Fab), the antibody heavy chain was successfully labeled with TAMRA-Q by pG(Fab)-EzMTG (**Fig. 2a, S2**). As the Lys and Gln residues in the pG(Fab) used in the present study were substituted with Arg and Asn residues, respectively, self-labeling of the pG(Fab)-EzMTG was not detected.

**Fig. 2.**
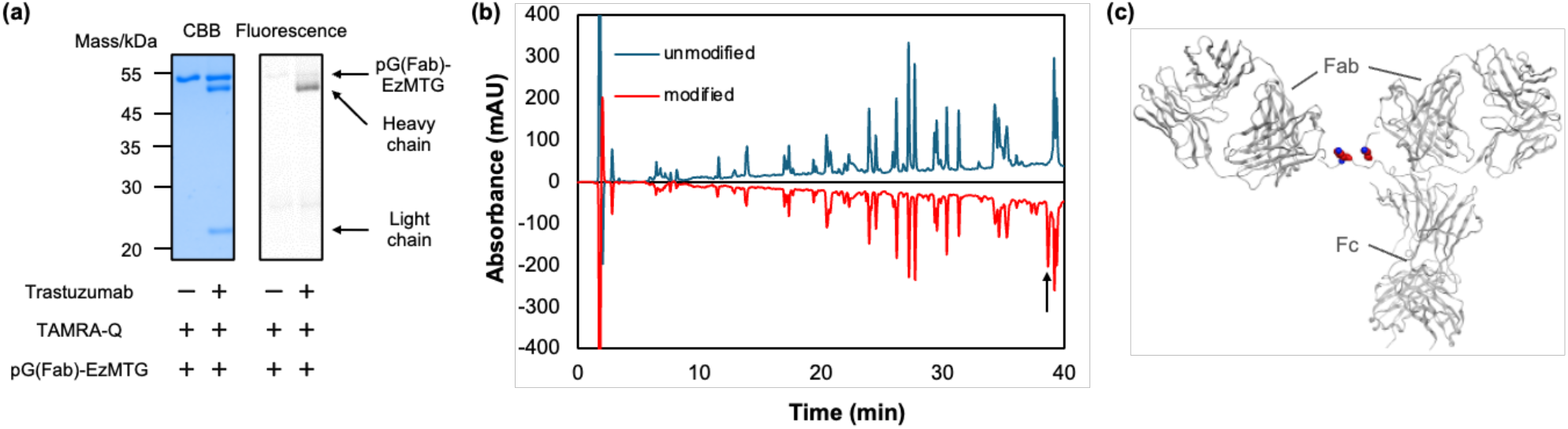
(a) Evaluation of the crosslinking activity of pG(Fab)-EzMTG using SDS-PAGE gel analysis and fluorescent imaging. The reaction was conducted with trastuzumab (1.3 μM), pG(Fab)-EzMTG (2.6 μM), and TAMRA-Q (100 μM) in 40 mM Tris-HCl (pH 8.0) at 37 °C for 60 min. Raw images of electrophoretic gels (CBB staining, left) and the fluorescence images (derived from TAMRA, right) are shown in Fig. S2 (ESI†). (b) Identification of Lys residues modified with TAMRA-Q by pG(Fab)-EzMTG. RP-HPLC chromatographs of trastuzumab modified by pG(Fab)-EzMTG. (c) Three-dimensional structure of trastuzumab. Trastuzumab is shown in gray. Lys225 of trastuzumab is represented as a space-filling model. The PDB entries used for Fab and Fc are 1N8Z.pdb and 1N8Z.pdb, respectively.

Next, peptide mapping analysis was performed on the TAMRA-Q-modified trastuzumab prepared using pG(Fab)-EzMTG, to identify the conjugation site. Interestingly, in contrast to EzMTG-pG(Fab), in which Lys65 was preferentially modified, Lys225, which is located in the hinge region connecting the Fc and Fab heavy chains of trastuzumab, was modified almost exclusively by TAMRA-Q, although minor modification was also detected at Lys65 and Lys293 (**Fig. 2b, 2c, S3**, and **S4**). In other words, simply switching the orientation of the fusion construct was sufficient to change the main modification site from Lys65 to Lys225 without altering the catalytic domain of microbial transglutaminase. Notably, although Lys225 has been reported as one of several modification sites in trastuzumab,^31, 44^ modifications occurring predominantly and with high efficiency at Lys225 appears to be unprecedented, highlighting a key feature of pG(Fab)-EzMTG.

**Fig. 3.**
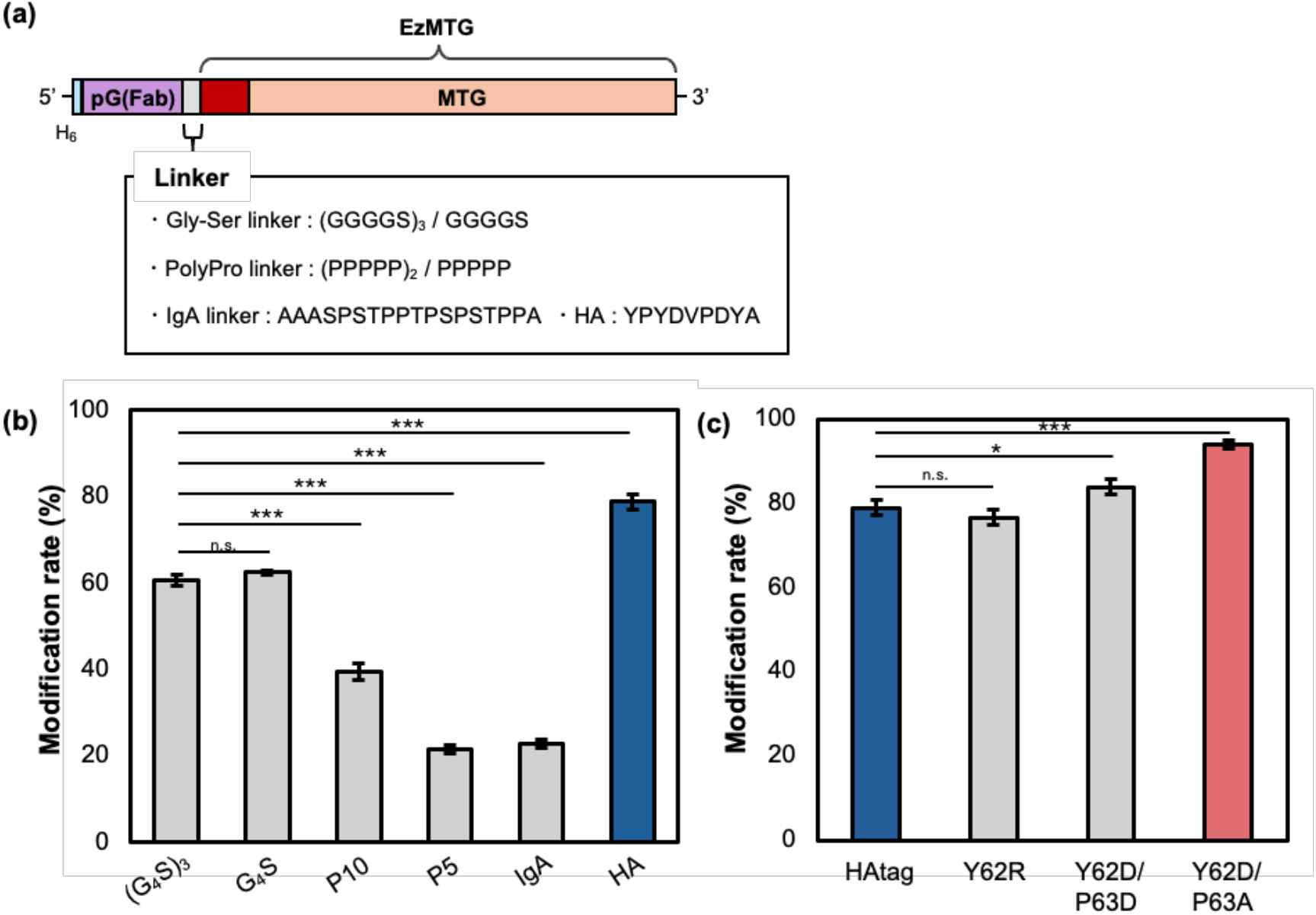
(a) Schematic representation of pG(Fab)-EzMTG variants with different linker sequences designed in the present study. Modification efficiency of trastuzumab by (b) pG(Fab)-EzMTG mutants and (c) pG(Fab)-HA-EzMTG mutants. The reaction was conducted with trastuzumab (1.3 μM), pG-fused EzMTG mutant (2.6 μM), and TAMRA-Q (100 μM) in 40 mM Tris-HCl (pH 8.0) at 37 °C for 120 min. *n* = 3; mean ± SE; \**p* < 0.05, \*\*\**p* < 0.001; n.s., not significant.

At least two factors may account for the selective modification of Lys225. First, the structural configuration of pG alters the set of Lys residues that EzMTG can access. As illustrated in **Fig. 1**, fusing EzMTG to the *N*- or *C*-terminus of pG(Fab) leads to obviously different spatial positioning of EzMTG in relation to the antibody. In EzMTG-pG(Fab), EzMTG is positioned near the antigen-binding region; however, in pG(Fab)-EzMTG, EzMTG is positioned closer to the hinge region. To better understand the orientation of EzMTG, in silico calculations of the encounter frequencies of the catalytic center of EzMTG were conducted with each pG-fused EzMTG and Fab complex (**Fig. S5**). The simulation analysis was performed with the Fab region without the hinge region. The results clearly showed that EzMTG-pG(Fab) showed higher encounter frequencies with Lys residues of the heavy chain located near the antigen-binding region (Lys30, Lys65, and Lys76), and Lys65 and Lys76 were among the residues predicted to be the most accessible. By contrast, pG(Fab)-EzMTG showed a distinct preference for Lys esidues close to the hinge region (Lys183 and Lys188 in the light chain and Lys136 in the heavy chain) (**Table S2**). Second, Lys225 exhibited a high positive residue patch area (PRPA) value among the Lys residues in the heavy chain of trastuzumab, with the second highest PRPA value (**Table S3**). These results showed a similar trend with Lys65 showing the highest PRPA value of all the Lys residues, and this residue was exclusively modified by EzMTG-pG(Fab), whereas the accessible Lys76 was not modified. Lys residues in positively charged regions were preferentially recognized by the negatively charged active site of EzMTG. Collectively, alteration of the orientation of EzMTG toward the antibody by pG, and the intrinsic electrostatic interaction, allowed control of the pG(Fab)-EzMTG modification site, resulting in preferential modification of Lys225.

### Optimization of the linker in pG(Fab)-EzMTG

Although Lys225 is located in the conserved hinge region, which can be applicable to a range of IgG antibodies, the modification rate using pG(Fab)-EzMTG was approximately 60% (**Fig. S6a, 6b**). This rate contrasts with the high labeling efficiency (over 90%) observed with EzMTG-pG(Fab).^42^ This reduced labeling efficiency is likely because of the more congested environment around Lys225 compared with Lys65, which may prevent EzMTG from accessing the target residue. To improve the modification rate, we focused on the linker connecting pG(Fab) and EzMTG. Flexible or rigid linkers are generally employed in fusion protein design, and appropriate linker selection is known to enhance the expression, activity, and stability of fusion proteins.^45^ We constructed a series of fusion proteins having different types of linkers of varying length, flexibility, and sequence between pG(Fab) and EzMTG. The modification efficiency of pG(Fab)-EzMTG mutants with flexible (Gly-Ser), rigid (polyPro), IgA^46^ and HA^47^ were evaluated (**Fig. 3a**).

The modification efficiency of most of these fusion proteins was lower, at approximately 20%–40%, compared with the Gly-Ser linker used in the original construct. However, the insertion of a HA linker markedly increased the labeling efficiency up to 80% (**Fig. 3b**). The HA linker consists of the peptide sequence, YPYDVPDYA, which contains aspartic acid (Asp, D) residues and therefore carries a net negative charge. The *N*-terminal sequence of the hinge site around the reactive Lys225 (…DKKVEPKSCDK_225_…) is enriched in positively charged Lys residues, thus electrostatic interactions between these Lys residues and Asp residues in the HA linker may facilitate closer proximity between EzMTG and Lys225. To examine the contribution of electrostatic interactions, we conducted the labeling reaction in Tris-HCl buffer containing 0.15 M NaCl and found that the modification efficiency decreased to approximately 50%, which was comparable to the efficiency observed with the (G₄S)₃ linker (**Fig. S6c**). These results support the concept that electrostatic interaction between the hinge region and HA linker contributed to enhancing the modification efficiency. These observations further suggest that the linker not only connects the two domains but also influences the local spatial relationship between the catalytic domain and the target residue, which may affect the modification efficiency.

We then attempted to optimize the sequence of HA linker to increase the labeling efficiency. Specifically, we targeted the two *N*-terminal residues of HA linker (Y_62_P_63_YDVPDYA; Y62 and P63). To introduce an additional negative charge, we substituted both residues with Asp, generating pG(Fab)-HA(Y62D/P63D)-EzMTG. This construct exhibited an increased modification rate of approximately 84% (**Fig. 3c**). In contrast, introducing a positively charged Arg into the HA linker, HA(Y62R), slightly reduced the modification rate compared with the original HA linker, although this difference was not statistically significant (**Fig. 3c**). These results further support the contribution of electrostatic interactions between the HA linker and the hinge site toward the modification efficiency, suggesting that Y62 is critical to position the EzMTG domain in appropriate orientation to the hinge site. We further designed pG(Fab)-HA(Y62D/P63A)-EzMTG where P63 was replaced with Ala to investigate the effect of proline. This variant exhibited even higher modification efficiency than pG(Fab)-HA(Y62D/P63D)-EzMTG (**Fig. 3c**) and showed higher modification efficiency relative to that of pG(Fab)-HA-EzMTG under identical NaCl-containing conditions (**Fig. S6d**). These results suggested that P63 plays a minimal role in electrostatic interactions between the HA linker and the region surrounding Lys225, instead contributing to modulating linker flexibility. pG(Fab)-HA(Y62D/P63A)-EzMTG, which showed the highest modification efficiency, was used in subsequent experiments.

### Evaluation of versatility of Lys225 modification

Having achieved a sufficient modification rate with trastuzumab, we next evaluated the general applicability of this system to different antibody species. The lack of generality is a limitation of the Lys65 modification by EzMTG-pG(Fab). Lys225, located in the hinge site of the constant region of IgG1 antibodies, is conserved in all the IgG antibodies investigated (trastuzumab, rituximab, and ramucirumab). Consequently, a high modification efficiency of approximately 95% was achieved for ramucirumab, which was comparable to that observed for trastuzumab. In contrast, the modification efficiency for rituximab was lower at approximately 70% (**Fig. 4a**). Despite the amino acid sequences surrounding Lys225 being nearly identical in the three antibodies, reduced efficiency was observed only for rituximab. We speculated that differences in the antibody-binding affinity of pG(Fab) may underlie the reduced labeling efficiency observed for rituximab. The binding affinity of pG(Fab) toward each IgG1 antibody was thus examined using biolayer interferometry (BLI). The results revealed that pG(Fab) exhibited a high binding ability in the order ramucirumab (*K*_D_: 28.2 nM) > trastuzumab (*K*_D_: 77.5 nM) > rituximab (*K*_D_: 318.7 nM) (**Fig. 4b**). As the reaction conditions used in the present study were optimized for trastuzumab, the weaker binding ability reduced the proximity effect, which is likely to account for the lower modification efficiency observed for rituximab. Increasing the concentration of pG(Fab)-HA(Y62D/P63A)-EzMTG increased the modification efficiency when the reaction was performed at an antibody-to-enzyme ratio of 1:4 (rituximab: MTG) (**Fig. 4a**). These results collectively indicated that, while the reactivity of pG(Fab)-EzMTG is influenced by its binding affinity toward individual IgG1 antibodies, optimizing the reaction conditions can achieve high modification efficiency across different antibodies. Therefore, pG(Fab)-EzMTG-mediated selective labeling of conserved Lys225 can be a versatile and broadly applicable strategy.

**Fig. 4.**
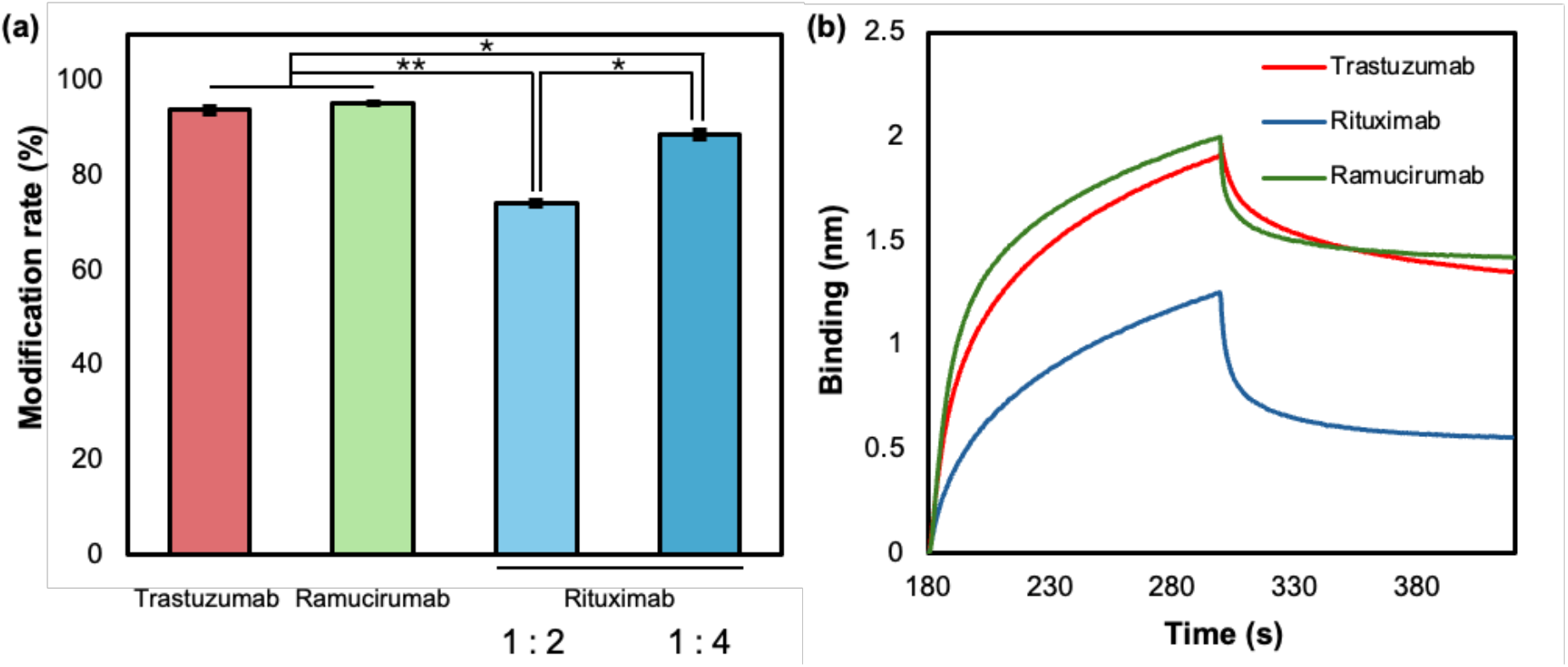
(a) Modification efficiency of pG(Fab)-HA(Y62D/P63A)-EzMTG with different IgG1 antibodies. The reaction was conducted with each antibody (1.3 μM), pG(Fab)-HA(Y62D/P63A)-EzMTG (2.6 μM), and TAMRA-Q (100 μM) in 40 mM Tris-HCl (pH 8.0) at 37 °C for 120 min. The reaction was also performed with rituximab (1.3 μM), pG(Fab)-HA(Y62D/P63A)-EzMTG (5.2 μM), and TAMRA-Q (200 μM) (antibody-to-enzyme ratio of 1:4). *n* = 3; mean ± SE; \**p* < 0.05, \*\**p* < 0.01. (b) Representative sensorgrams for evaluation of the binding affinity of pG(Fab)-HA(Y62D/P63A)-EzMTG to IgG1 antibodies using bio layer interferometry (BLI) analysis.

### Evaluation of antigen-binding ability of Lys225-modified trastuzumab

We evaluated the antigen-binding ability of TAMRA-Q-modified trastuzumab by assessing the binding to human epidermal growth factor receptor type 2 (HER2), which is the target antigen of trastuzumab. Trastuzumab modified with TAMRA-Q at Lys225 was purified from the crosslinked reaction mixture by separating pG(Fab)-EzMTG. Dissociation of pG(Fab)-EzMTG was achieved by cation-exchange chromatography under acidic conditions (pH 3.0), yielding TAMRA-Q-modified trastuzumab at approximately 76% purity (**Fig. S7**). The binding affinity of the purified TAMRA-Q-modified trastuzumab toward HER2 was then determined. The dissociation constant (*K*_D_) of the purified sample was 0.20 nM, demonstrating that the Lys225-modified trastuzumab retained a binding affinity comparable to that of the unmodified trastuzumab (*K*_D_ = 1.65 nM) (**Fig. 5a, Table S4**). Although the underlying reason for the slight increase in the apparent binding affinity of TAMRA-Q-modified trastuzumab, relative to the unmodified antibody, is not yet clear, a potential interaction between TAMRA-Q and HER2 cannot be excluded. Next, we evaluated the binding ability of TAMRA-Q-modified trastuzumab to HER2 expressed on the cell surface. The TAMRA-Q-modified trastuzumab was incubated with both HER2-positive SK-BR-3 cells and HER2-negative MDA-MB-231 cells, after which membrane binding was evaluated. Using confocal laser scanning microscopy (CLSM), we observed red fluorescence localized to the plasma membrane of SK-BR-3 cells following treatment with TAMRA-Q-modified trastuzumab. In contrast, no appreciable fluorescence signal was detected in the MDA-MB-231 cells. These results suggest that Lys225-modified trastuzumab retains intrinsic antigen specificity and can selectively target HER2-positive cells (**Fig. 5c, S8**).

**Fig. 5.**
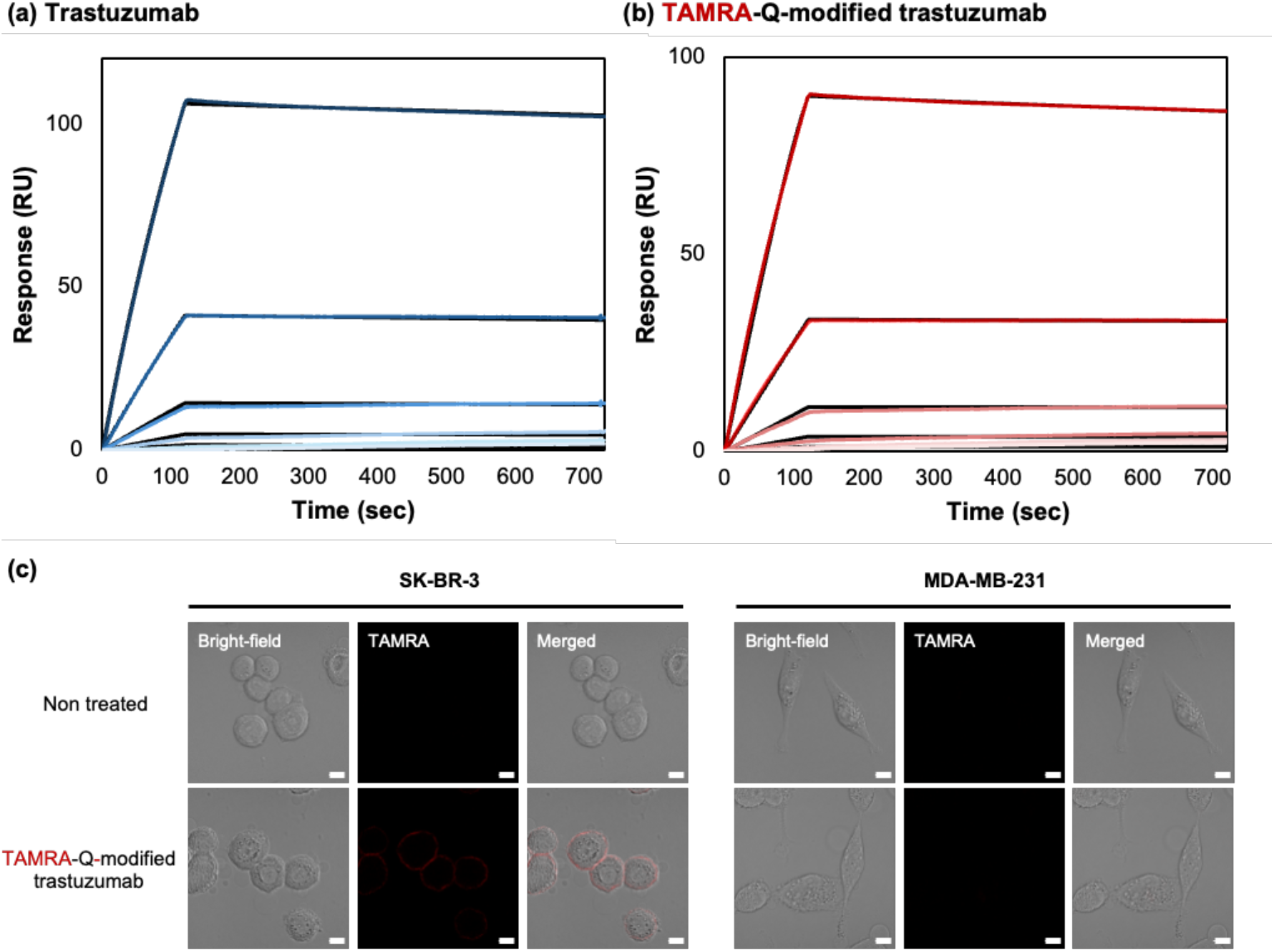
Binding of TAMRA-Q-modified trastuzumab to HER2. Sensorgrams of SPR analysis of the binding of HER2 at 0.494, 1.48, 4.44, 13.3, 40, and 120 nM against (a) trastuzumab and (b) TAMRA-Q-modified trastuzumab immobilized in the surface of a protein A chip. Raw data are colored blue in (a) and red in (b), whereas the fitted data are shown in black in both panels. (c) Representative fluorescence images obtained by CLSM. SK-BR-3 and MDA-MB-231 cells were incubated with each sample at 37 °C for 1 h. Bars: 10 μm.

### Preparation of ADCs and evaluation of cytotoxicity

Finally, we prepared an ADC by conjugating a modified Gln substrate (MMAE-PAB-vc-CYPLQMRG-NH₂, MMAE-Q; **Fig. S1**) bearing the potent cytotoxic payload monomethyl auristatin E (MMAE) to trastuzumab using pG(Fab)-HA(Y62D/P63A)-EzMTG. We analyzed the conjugation site in the same way as for the TAMRA-Q-modified antibody and determined that MMAE-Q was attached at Lys225 (**Fig. S9**). The resulting MMAE-Q-modified trastuzumab (ADC) was purified by cation-exchange chromatography under acidic conditions (pH 3.0), analogous to the purification procedure for TAMRA-Q-modified trastuzumab, affording purified ADC with a modification efficiency of approximately 80% (**Fig. S10a, b**). The ADC constructed in this way exhibited an average DAR value of 1.83 (**Fig. S10c**). Although the DAR = 2 species was the most abundant peak, minor populations corresponding to DAR = 3–4 were also detected. These results suggest that, while MMAE-Q was primarily conjugated to Lys225, a small fraction of the payload was additionally attached to other Lys residues, likely attributable to minor additional conjugation at Lys65 and Lys293.

The cytotoxic activity of the constructed ADC was evaluated. Treating HER2-positive SK-BR-3 cells with the ADC resulted in markedly enhanced potency, with a half maximal inhibitory concentration (IC₅₀) value of 43 pM, compared with trastuzumab alone (**Fig. 6a**). This activity was comparable to that of previously reported trastuzumab-MMAE conjugates (IC₅₀ = 21.7^22^ and 120^24^ pM), as well as the FDA-approved ADC Kadcyla^®^ (approximately 0.01 ug/mL, corresponding to ca. 67 pM^48^). These results support the potential therapeutic efficacy of the present construct. By contrast, minimal cytotoxicity was observed in HER2-negative MDA-MB-231 cells treated with MMAE–trastuzumab (Lys225) (**Fig. 6b**), indicating that the cytotoxic effect was antigen-dependent. These results demonstrate that Lys225-selective modification can be applied to the preparation of ADC while retaining the antigen-dependent activity of trastuzumab.

**Fig. 6.**
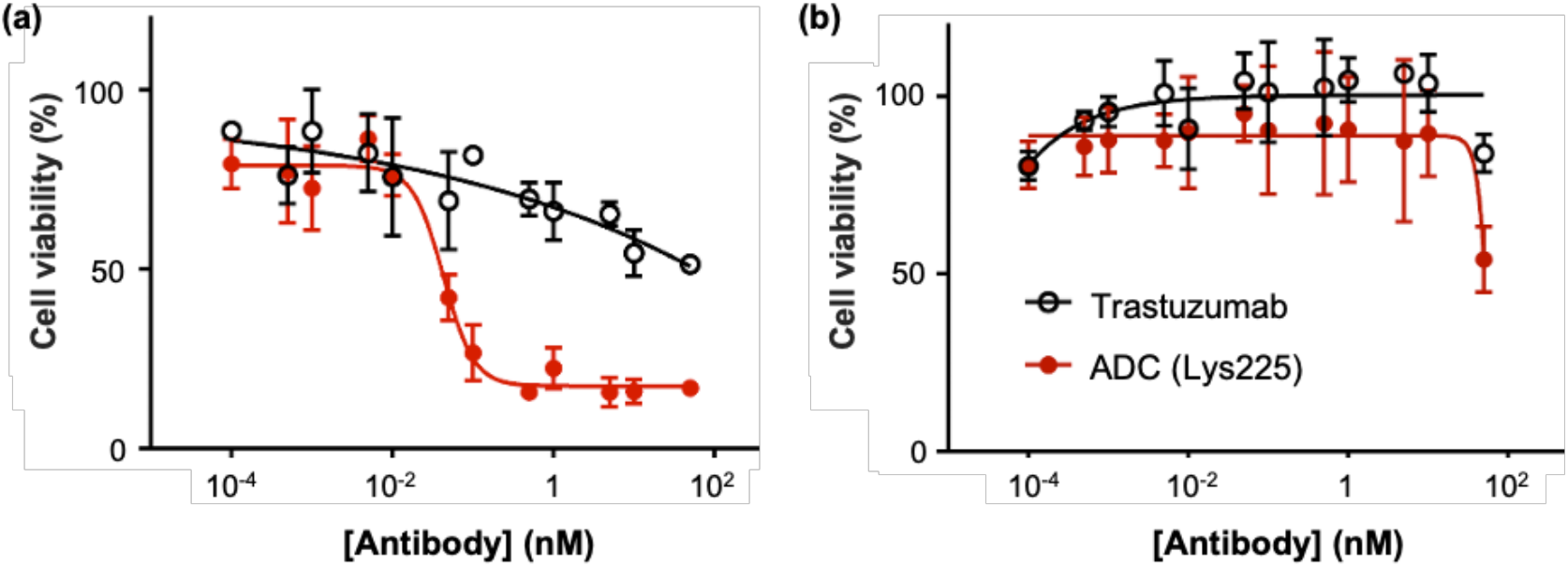
Cytotoxicity assays with (a) SK-BR-3 cells (HER2+) and (b) MDA-MB-231 cells (HER2–). All assays were performed in triplicate.

## Conclusion

By fusing an antibody-binding protein [pG(Fab)] to the activated precursor of MTG (EzMTG), we exploited a proximity effect to achieve site-selective modification of Lys225 in the heavy chain of native IgG1 antibodies. The results demonstrated that the modification site can be dramatically altered through the design of the fusion protein. Enhanced modification rates were observed for three different commercially available IgGs (trastuzumab, rituximab, and ramucirumab) by optimizing the linker connecting the fusion proteins, which showed the critical role of peptide linkers in the fusion protein design. As an application of this approach, an ADC prepared using the optimized fusion protein, pG(Fab)-HA(Y62D/P63A)-EzMTG, exhibited an average DAR of 1.83 and antigen-dependent cytotoxicity. These findings provide a powerful strategy for controlling antibody modification sites by engineering fusion protein partners. Collectively, this work introduces a generalizable strategy for controlling enzymatic selectivity via proximity effects, providing a new conceptual framework for protein modification and bioconjugation.

## Methods

### Materials

Miller lysogeny broth (LB), tryptone, dried yeast extract, tris(hydroxymethyl)aminomethane (Tris), guanidine hydrochloride, L-(+)-arginine monohydrochloride, and hydrochloric acid were purchased from Nacalai Tesque, Inc. (Kyoto, Japan). Potassium dihydrogen phosphate, dipotassium hydrogen phosphate, disodium hydrogen phosphate dodecahydrate, sodium chloride, dithiothreitol, ammonium bicarbonate, trehalose, 4% paraformaldehyde phosphate buffer solution, acetonitrile (ACN), and sodium ampicillin were purchased from Fujifilm Wako Pure Chemical Corporation (Osaka, Japan). Glycerol, citric acid, tri-sodium citrate, and potassium chloride were purchased from Kishida Chemical Co., Ltd (Osaka, Japan). Tween 20 was purchased from Tokyo Chemical Industry (Tokyo, Japan). Trifluoroacetic acid (TFA) was purchased from Watanabe Chemical Industry (Hiroshima, Japan). Human IgG1 trastuzumab (Herceptin^®^) and chimeric IgG1 rituximab (MabThera^®^) were purchased from F. Hoffmann-La Roche, Ltd. (Basel, Switzerland). Human IgG1 ramucirumab was purchased from Eli Lilly Japan K.K. (Kobe, Japan). Imidazole, trypsin, α-cyano-4-hydroxycinnamic acid (CHCA), TAMRA-YPLQMRG-NH_2_, and Ac-CYPLQMRG-NH_2_ were purchased from Sigma-Aldrich Co. LLC (St. Louis, MO, USA). MC-Val-Cit-PAB-MMAE was purchased from BroadPharm (San Diego, CA, USA). SK-BR-3 (ATCC HTB-30) and MDA-MB-231 (ATCC HTB-26) cells were obtained from the American Type Culture Collection (Manassas, VA, USA). McCoy’s 5A medium and Dulbecco’s modified Eagle medium (DMEM) were purchased from Thermo Fisher Scientific (Waltham, MA, USA). The protein structures were generated by Molecular Operating Environment (Chemical Computing Group Inc., Quebec, Canada).

### Construction of pG-fused EzMTG mutants

The DNA sequence coding the pG(Fab) sequence was amplified using PCR, and the product was inserted into a pET22b+ vector carrying the gene coding engineered zymogen of microbial transglutaminase (EzMTG) using an In-Fusion HD cloning kit (Takara Bio Inc., Shiga, Japan). The DNA sequences coding the pG(Fab) sequences were amplified using PCR. The products were inserted into the pET22b+ vector carrying the gene encoding EzMTG using the In-Fusion HD cloning kit. The amino acid sequences of all the proteins used in the present study are summarized in Table S1.

### Expression and purification of pG(Fab)-EzMTG mutants

The expression of the pG(Fab)-EzMTG mutants was conducted using *Escherichia coli* BL21 star (DE3). The plasmid vectors encoding pG(Fab)-EzMTG mutants were transformed into cells by heat shock and cultured on an LB plate containing 100 μg/mL sodium ampicillin. A single colony was inoculated into 5 mL of LB containing the same amount of sodium ampicillin and shaken at 220 rpm at 37 °C for 6 h. A preculture of cells expressing pG-fused EzMTG was placed into 1 L of Terrific broth medium containing 100 μg/mL ampicillin and cultured with shaking at 220 rpm at 37 °C until the Optical Density at 600 nm (OD_600_) value reached 0.4–0.5. The expression of pG(Fab)-EzMTG mutants was induced by the addition of 0.1 mM isopropyl-β-D-thiogalactoside followed by shaking at 18 °C. pG(Fab)-EzMTG mutants were cultured in an auto-induction medium. The cells were harvested by centrifugation at 5,000 ×*g* for 20 min, and the supernatant was discarded. The pellet was resuspended in phosphate-buffered saline (PBS, pH 7.4) and frozen at −80 °C until protein purification.

For the purification of pG(Fab)-EzMTG mutants, the cell pellet was thawed in running water. The cell suspension was sonicated on ice for 12.5 min. The cell debris was removed by centrifugation at 15,000 ×*g* for 60 min at 4 °C. The supernatants were initially applied to a crude column containing 5 mL of HisTrap FF (Cytiva, Tokyo, Japan), which was pre-equilibrated with 93% HisTrap binding buffer (20 mM Tris-HCl, 500 mM NaCl, pH 7.4) and 7% HisTrap elution buffer (20 mM Tris-HCl, 500 mM NaCl, 500 mM imidazole, pH 7.4). The column was washed with the binding buffer until all the unbound substances had been eluted out; then, the proteins were eluted with an elution buffer with a gradient of 7% to 100%. Each pG(Fab)-EzMTG mutant was purified by size exclusion chromatography using HiLoad 16/600 Superdex 200 pg (Cytiva, Tokyo, Japan) with 1×PBS with 0.1 mM dithiothreitol (pH 7.4) as a running buffer. The purified pG(Fab)-EzMTG mutants were concentrated using 30 kDa molecular weight cutoff (MWCO) Amicon Ultra-15 centrifugal filter units (Millipore, Tokyo, Japan). To determine protein concentrations, the absorbances at 280 nm were measured using a NanoDrop 2000c spectrophotometer (Thermo Fisher Scientific).

### Evaluation of the crosslinking activity of pG(Fab)-EzMTG mutants

Trastuzumab (1.3 μM) was mixed with each pG(Fab)-EzMTG mutant (2.6 μM) in 40 mM Tris-HCl (pH 8.0). The crosslinking reaction was started by the addition of TAMRA-Q (100 μM, total volume: 50 μL). The reaction was performed at 37 °C for 60 min. A 2× sodium dodecyl sulfate–polyacrylamide gel electrophoresis (SDS-PAGE) sample buffer was added to the reaction solution, and the mixed solution was heated at 98 °C for 2 min. After denaturation treatment, SDS-PAGE was conducted using 12.5% acrylamide gel with Coomassie brilliant blue staining (Quick-CBB, Fujifilm Wako Pure Chemical Corporation). The progression of substrate modification was determined using iBright FL1500 (Thermo Fisher Scientific).

### Identification of modification position

Unmodified trastuzumab and TAMRA-Q-modified trastuzumab were fragmented by reduction-alkylation and trypsin treatment. The resulting peptide fragments were injected into a COSMOSIL 5C18-AR-300 (4.6 ID × 150 mm) column for RP-HPLC analysis. In the RP-HPLC, water and ACN containing 0.1% TFA were used, and the analysis conditions were as follows: flow rate, 1.2 mL min^-1^; detection wavelengths, 214, 280, and 565 nm; and gradient, water: ACN (0.1% TFA) = 100:0 to 60:40 over 40 min. The molecular weight of each peak-derived sample fractionated was determined by MALDI-TOF-MS. Sample solutions were spotted onto a CHCA matrix and mass spectra were obtained using a Bruker Autoflex max MALDI-TOF mass spectrometer (Bruker, Billerica, MA, USA) in positive ion detection mode. The position of modification was identified by calculating the mass of the peptide fragment from the obtained mass using the following formula and comparing it with the theoretical mass (SCDK_225_THTCPPCPAPELLGGPSVFLFPPKPK = 3337.594). Theoretical masses were calculated using Expasy PeptideMass (Swiss Institute of Bioinformatics, Lausanne, Switzerland). *N*-terminal analysis of the modified antibody fragment was contracted to the Japan Institute of Leather Research (Ibaraki, Japan).

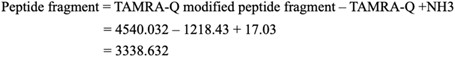

### Modeling and simulation of pG-fused EzMTG

Structure models of pG-fused EzMTG were first constructed by homology modeling. Initial structures were constructed by simply joining the pG-Fab part (Protein Data Bank (PDB) ID: 1IGC) [doi:10.1006/jmbi.1994.1691] and the MTG part (PDB ID: 3IU0) by the MODELLER [doi:10.1006/jmbi.1993.1626] software. Both EzMTG-pG(Fab) and pG(Fab)-EzMTG variants were modeled and used in the subsequent analysis. After constructing the initial structures, coarse-grained simulations were performed to investigate whether EzMTG’s reaction center can geometrically access Lys(K) residues (details described later). We used AICG2+ [doi:10.1073/pnas.1402768111] coarse-grained potential with the Go-like potential applied within pG, Fab, and EzMTG, and between pG-Fab. The flexible linker part was modeled by the flexible local potential [doi:10.1016/j.bpj.2011.08.003]. Simulation was performed with CafeMol[doi:10.1021/ct2001045], and the simulation was performed for 10^7^ steps with the “tstep” value of 0.1. The initial 2×10^6^ steps were discarded as the equilibration. 240 independent simulations were performed, each with a different random number seed, for each initial model (Fig. S5(a)).

We defined “pre-reaction collision event” as a Cα atom of a Lys residue being within 20 Å of Cys 114 (reaction center) Cα atom in EzMTG. Using the definition, we calculated the event ratio (the number of event divided by total trajectory snapshots) for each Lys residue. The ratio was then back-mapped on the structure as presented in Fig. S5(b). Figures were created using PyMOL [The PyMOL Molecular Graphics System, Version 2.5.0, Schrödinger, LLC.] and VMD [doi:10.1016/0263-7855(96)00018-5].

### Evaluation of modification rate

Trastuzumab (1.3 μM) was mixed with each MTG mutant (2.6 μM) in 40 mM Tris-HCl (pH 8.0) with 2 v/v% trehalose. The crosslinking reaction was started by adding TAMRA-Q (100 μM, total volume: 50 μL). The reaction was performed at 37 °C for 120 min. The reaction solution was sampled using 30 μL aliquots, which were mixed with 72 μL of 6 M guanidine hydrochloride containing 100 mM ammonium bicarbonate and 8 μL of 1 M dithiothreitol. The reduction reaction was then carried out at 60 °C for 30 min. Denatured and reduced samples (100 μL) were injected into a COSMOSIL 5Ph-AR-300 (4.6 ID × 150 mm) column for RP-HPLC analysis. In the RP-HPLC, water and ACN containing 0.1% TFA were used, and the analysis conditions were: flow rate, 1.2 mL min^-1^; detection wavelength, 214 nm; gradient, water: ACN (0.1% TFA) = 100:0 to 72:28 over 2 min then 72:28 to 62:38 over 16 min. The modification rate was calculated from the following formula using the peak areas of the TAMRA-Q-modified and unmodified heavy chains.

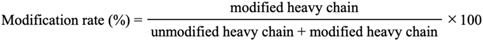

### Evaluation of the antibody-binding ability of pG(Fab)-EzMTG mutant using bio-layer interferometry analysis with BLItz

Bio-layer interferometry was performed with a BLItz (Fortebio, CA, USA) to analyze the binding of each pG(Fab)-HA(Y62D/P63A)-EzMTG to each IgG1 antibody (trastuzumab, rituximab, and ramucirumab). Streptavidin (SA) biosensors (Sartorius AG, Göttingen, Germany) were hydrated in Milli-Q water for at least 10 min before use. To immobilize each IgG1 antibody on the SA biosensors, each IgG1 antibody was biotinylated using a Biotin Labeling Kit-NH_2_ (Dojindo Laboratories, Kumamoto, Japan). After the baseline step with kinetics buffer (Fortebio, CA, USA), 4 μL of each biotinylated IgG1 antibody (30 μg/mL in Milli-Q water) was added to the drop holder to allow the association of biotin-IgG1 antibody with the SA biosensor surfaces for 120 s. After the baseline step with kinetics buffer for 30 s, the association of samples (0.125, 0.25, 0.50, and 1.0 μM of pG(Fab)-EzMTG) with the proteins on the tips for 120 s, and a dissociation step for 120 s by soaking in kinetics buffer, were performed.

### Purification of TAMRA-Q-modified trastuzumab

Trastuzumab (2.6 μM) was mixed with pG(Fab)-HA(Y62D/P63A)-EzMTG (5.2 μM) in 40 mM Tris-HCl (pH 8.0). The crosslinking reaction was started by adding TAMRA-Q (200 μM, total volume: 2 mL). The reaction was performed at 37 °C for 120 min. The MTG reaction was stopped and the antibody-binding ability of protein G was inactivated by adding 4 mL of 0.1 M citrate buffer (pH 3.0) to the reaction solution. The sample solution was injected into a Protein Ark HiFliQ S-type column for cation-exchange chromatography. In the cation-exchange chromatography, 50 mM citrate buffer A (0.2 M arginine hydrochloride, pH 3.0) and 50 mM citrate buffer B (0.5 M arginine hydrochloride, 2 M sodium chloride, pH 3.0) were used, and the analysis conditions were as follows: flow rate, 1.2 mL min^-1^; detection wavelengths, 280 and 565 nm; gradient, A: B = 95:5 to 45:55 over 20 min. Each fraction was neutralized by adding 100 µL of 1 M Tris-HCl (pH 8.0). Fraction 1 (**F1**) containing TAMRA-Q-modified trastuzumab and unmodified trastuzumab was collected to completely separate the antibody and pG(Fab)-EzMTG (Fig. S7a). Fraction 2 (**F2**) contained primarily pG(Fab)-HA(Y62D/P63A)-EzMTG and the residual TAMRA-Q-modified trastuzumab, suggesting the presence of a complex of TAMRA-Q-modified trastuzumab and pG(Fab)-HA(Y62D/P63A)-EzMTG that could not be separated under the experimental conditions. The fraction **F1** was concentrated, and the buffer was exchanged to 1× PBS (pH 7.4) using 10 kDa MWCO Amicon Ultra-15 centrifugal filter units. The purity of TAMRA-Q-modified trastuzumab in **F1** was determined from the RP-HPLC chromatograms (Fig. S7b).

### Evaluation of the antigen-binding ability of native and modified trastuzumab using surface plasmon resonance (SPR)

SPR measurements were performed on a Biacore™ 8K instrument (GE Healthcare, Uppsala, Sweden) to determine the kinetic parameters of trastuzumab and TAMRA-Q-modified trastuzumab binding to HER2 (Sino biological Inc., Beijing, China). Antibodies were captured on a Sensor Chip Protein A (Cytiva) in PBS-T (0.005% Tween 20) following the manufacturer’s instructions. HER2 was injected under multi-cycle kinetic conditions at 30 μL/min, with association and dissociation times of 120 and 600 s, respectively. The sensor surface was regenerated using 10 mM glycine (pH 1.5) for 30 s. All measurements were performed in PBS-T (0.005% Tween 20) at 25 °C. Analysis was conducted using the Biacore™ 8K evaluation software using a 1:1 binding kinetic fitting model. Biacore™ 8K measurements were performed at the Exploratory Research Center on Life and Living Systems (ExCELLS), National Institutes of Natural Sciences (NINS).

### Cell Culture

SK-BR-3 cells and MDA-MB-231 cells were purchased from RIKEN Cell Bank (Ibaraki, Japan). SK-BR-3 cells were cultured in McCoy’s 5A medium and MDA-MB-231 cells were cultured in DMEM medium (high glucose). The medium was supplemented with 10% fetal bovine serum and 1% antibiotic–antimycotic mixed solution, and cultured in a humidified incubator at 37 °C in the presence of 5% CO_2_.

### Confocal Laser Scanning Microscopy (CLSM) observations

Cells were plated in a multi-well glass-bottom dish (10000 cells per well) and allowed to attach to the glass-bottom dish for 24 h at 37 °C. After washing with Opti-MEM, TAMRA-Q-modified trastuzumab (final conc.: 10 μg/mL) was added and the cells were incubated at 37 °C for 1 h. After washing again with Opti-MEM, 100 µL of 4% paraformaldehyde phosphate buffer solution was added and the cells were allowed to stand for 10 min to fix the cells. The cells were observed using CLSM (Carl Zeiss microscope, Oberkochen, Germany) with diode lasers (567 nm for TAMRA).

### Purification of MMAE-Q-modified trastuzumab (ADC)

Trastuzumab (2.6 μM) was mixed with pG(Fab)-HA(Y62D/P63A)-EzMTG (5.2 μM) in 40 mM Tris-HCl (pH 8.6). The crosslinking reaction was started by adding MMAE-Q (400 μM, total volume: 2 mL). The reaction was performed at 37 °C for 120 min. The MTG reaction was stopped and the antibody-binding ability of protein G was inactivated by adding 4 mL of 0.1 M citrate buffer (pH 3.0) to the reaction solution. The sample solution was injected into a Protein Ark HiFliQ S-type column for cation-exchange chromatography. In the cation-exchange chromatography, 50 mM citrate buffer A (0.2 M arginine hydrochloride, pH 3.0) and 50 mM citrate buffer B (0.5 M arginine hydrochloride, 2 M sodium chloride, pH 3.0) were used, and the analysis conditions were as follows: flow rate, 1.2 mL min^-1^; detection wavelengths, 280 and 565 nm; gradient, A: B = 95:5 to 45:55 over 20 min. Each fraction was neutralized by adding 100 µL of 1 M Tris-HCl (pH 8.0). Fraction 1 (**F1**) containing MMAE-Q-modified trastuzumab and unmodified trastuzumab was collected to completely separate the antibody and pG(Fab)-HA(Y62D/P63A)-EzMTG (Fig. S11a). Fraction 2 (**F2**) contained primarily pG(Fab)-EzMTG and the residual MMAE-Q-modified trastuzumab, suggesting the presence of a complex of MMAE-Q-modified trastuzumab and pG(Fab)-HA(Y62D/P63A)-EzMTG that could not be separated under the experimental conditions. The fraction **F1** was concentrated, and the buffer was exchanged to 1× PBS (pH 7.4) using 10 kDa MWCO Amicon Ultra-15 centrifugal filter units. The purity of MMAE-Q-modified trastuzumab in **F1** was determined from the RP-HPLC chromatograms (Fig. S11b).

### Evaluation of cytotoxicity

For SK-BR-3 and MDA-MB-231 cell lines, the cells were plated into 96-well plates (at 5,000 cells per well), and the plates were incubated for 24 h at 37 °C with 5% CO_2_. SK-BR-3 cells were cultured in McCoy’s 5A medium and MDA-MB-231 cells were cultured in DMEM (high glucose). The medium was supplemented with 10% heat-inactivated fetal bovine serum and 1% antibiotic–antimycotic mixed solution. The ADC samples were diluted using the corresponding medium from 10 to 0.005 (0.01) nM and then added to the wells in triplicate for each concentration. The cells were cultured at 37 °C with 5% CO_2_ for 3 days, then cell counting kit-8 solution (Dojindo) was added. The absorbance of formazan released by viable cells was measured at 450 nm using a microplate reader after incubation at 37 °C with 5% CO_2_ for 4 h. Finally, the cell viability curve and IC_50_ values were calculated using GraphPad Prism software.

### Intact mass analysis

MMAE-Q-modified trastuzumab (10 μg) was dissolved in 100 μL of PBS. For deglycosylation, 4 units of PNGase F (Roche, Mannheim, Germany) were added in the sample solution, and the mixture was incubated at 37°C for overnight. Finally, 2 μL of the digested solution was directly injected into the LC-MS instrument. The LC-MS condition and data analysis were performed in accordance with our previous report^49^.

## Author information

### Author Contributions

The manuscript was written through contributions of all authors. R.N.: data curation, formal analysis, methodology, investigation, validation, writing the original draft; K.M., Y.K., M.K., S.S., N.H., A.S., J.M.M.C.: data curation, formal analysis, methodology, validation; M.U.: methodology, investigation; N.K.: conceptualization, supervision, methodology, validation, writing/reviewing, and editing. All authors have given approval to the final version of the manuscript.

## Acknowledgment

This study was supported by AMED, Japan (Grant Numbers: JP21ae0121003 to M.U., JP21ae0121004 to N.K., JP21ae0121005 to S.S., JP21ae0121006 and 22ak0101185 to N.H.), and JSPS KAKENHI (Grant number: JP23H00247 to N.K.). Victoria Muir, PhD, from Edanz (https://jp.edanz.com/ac) edited the English text of a draft of this manuscript. This research was partially supported by Joint Research of the Exploratory Research Center on Life and Living Systems (ExCELLS) (ExCELLS program No.25EXC321) and by grants from the Japan the Platform Project for Supporting Drug Discovery and Life Science Research (Basis for Supporting Innovative Drug Discovery and Life Science Research [BINDS]) from AMED (Grant No.25ama121031 to J.M.M.C.). We thank Ms. C. Obata from NIHS for sample preparation and LC/MS analysis. The simulation was carried out using the General Project category on supercomputer “Flow” at Information Technology Center, Nagoya University.

## Supporting Information

**Fig. S1.**
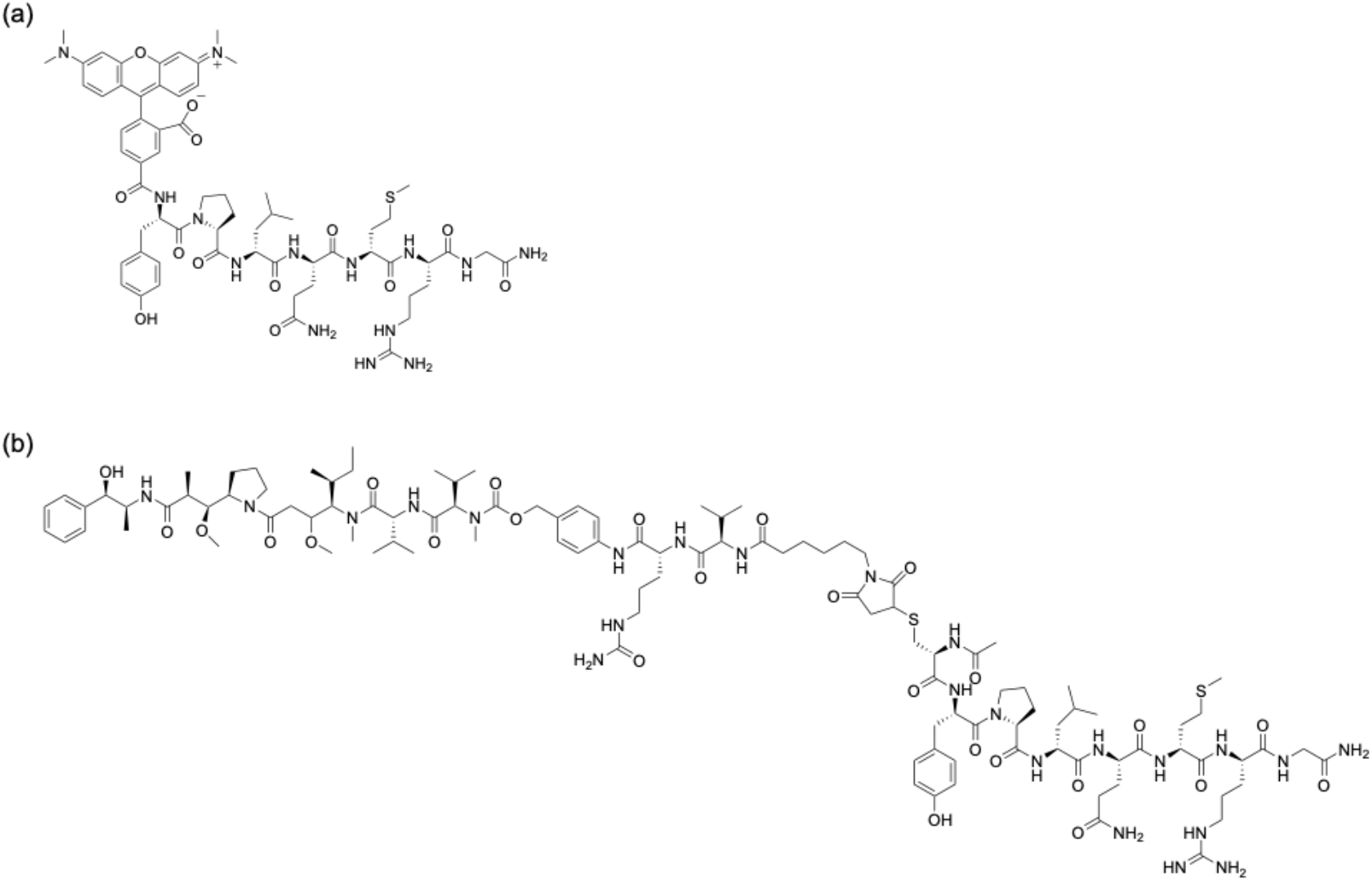
Chemical structures of (a) TAMRA-YPLQMRG-NH2 and (b) MMAE-PAB-vc-CYPLQMRG-NH2 (PAB, p-aminobenzyloxycarbonyl; vc, Val-Cit).

**Fig. S2.**
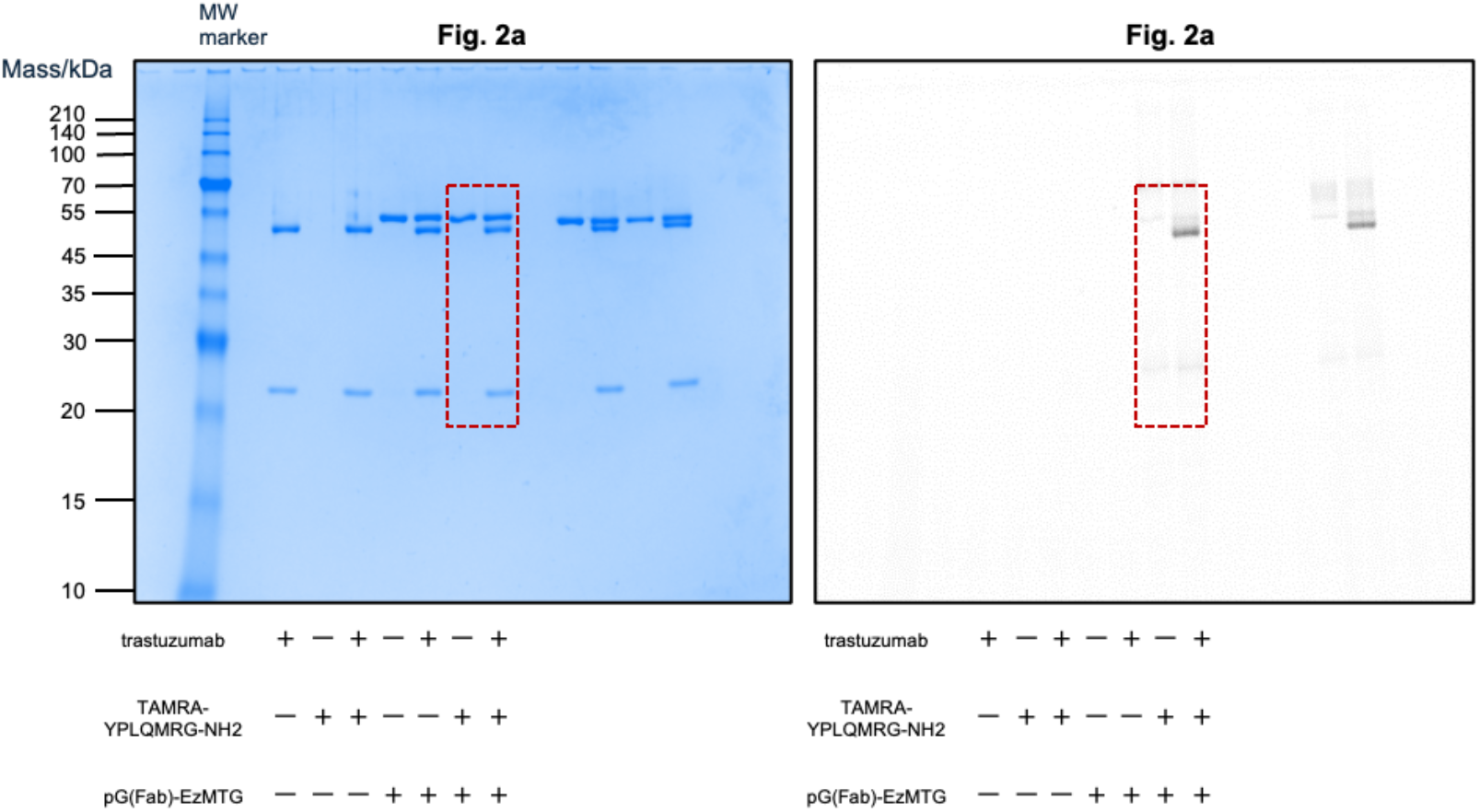
Raw images of sodium dodecyl sulfate–polyacrylamide gel electrophoretic gels (CBB staining, left) and the fluorescent images (derived from TAMRA, right) for the data presented in Fig. 2a (images in Fig. 2a are indicated by the red-dashed rectangular boxes).

**Fig. S3.**
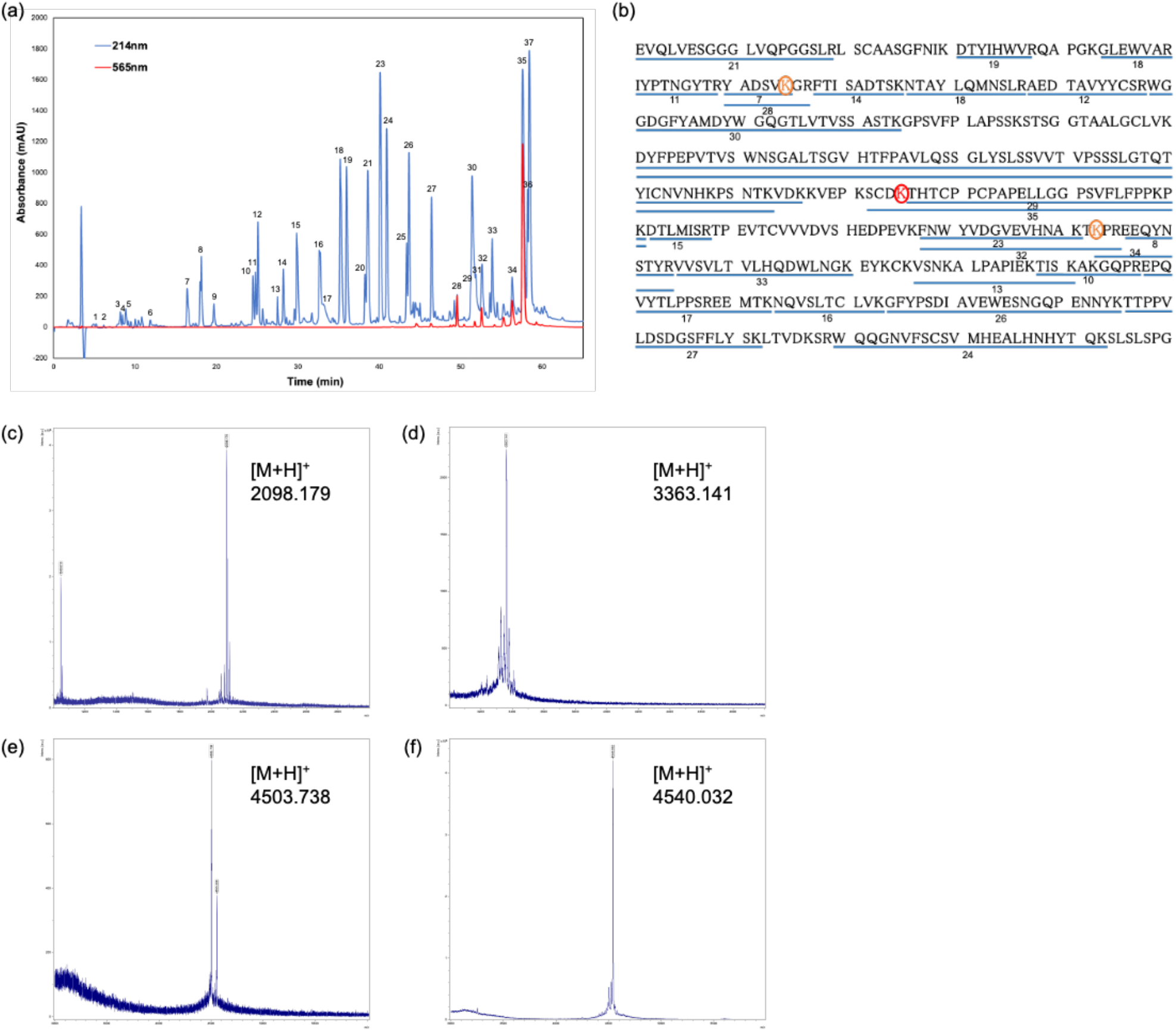
Identification of modification positions. (a) Chromatogram of TAMRA-Q-modified heavy chain fragmentation peptides and (b) peptide mapping results shown on the heavy chain sequence [the numbers at the bottom correspond to the peak numbers on the chromatogram in (a)]. Lys225 is shown in red, and Lys65 and Lys293 are shown in orange. MALDI-TOF-MS results for the fragments from peaks (c) No. 28, (d) No. 32, (e) No. 34, and (f) No. 35 in the chromatogram in (a).

**Fig. S4.**
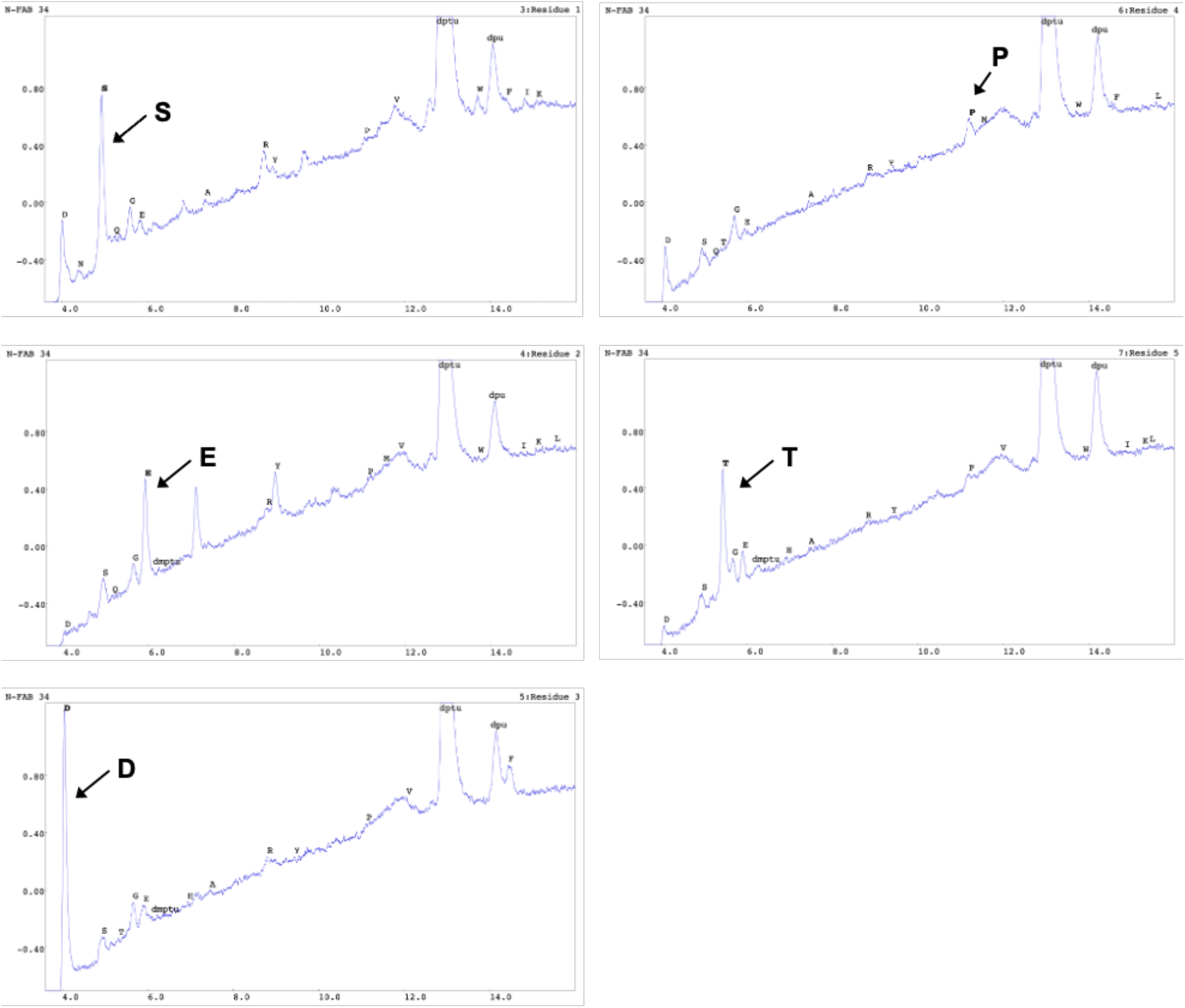
HPLC chromatograms of each residue obtained from *N*-terminal sequence analysis of modified antibody fragment. The results for the second and fourth residues were ambiguous, likely because the second residue corresponds to an alkylated Cys and the fourth residue to a chemically modified Lys.

**Fig. S5.**
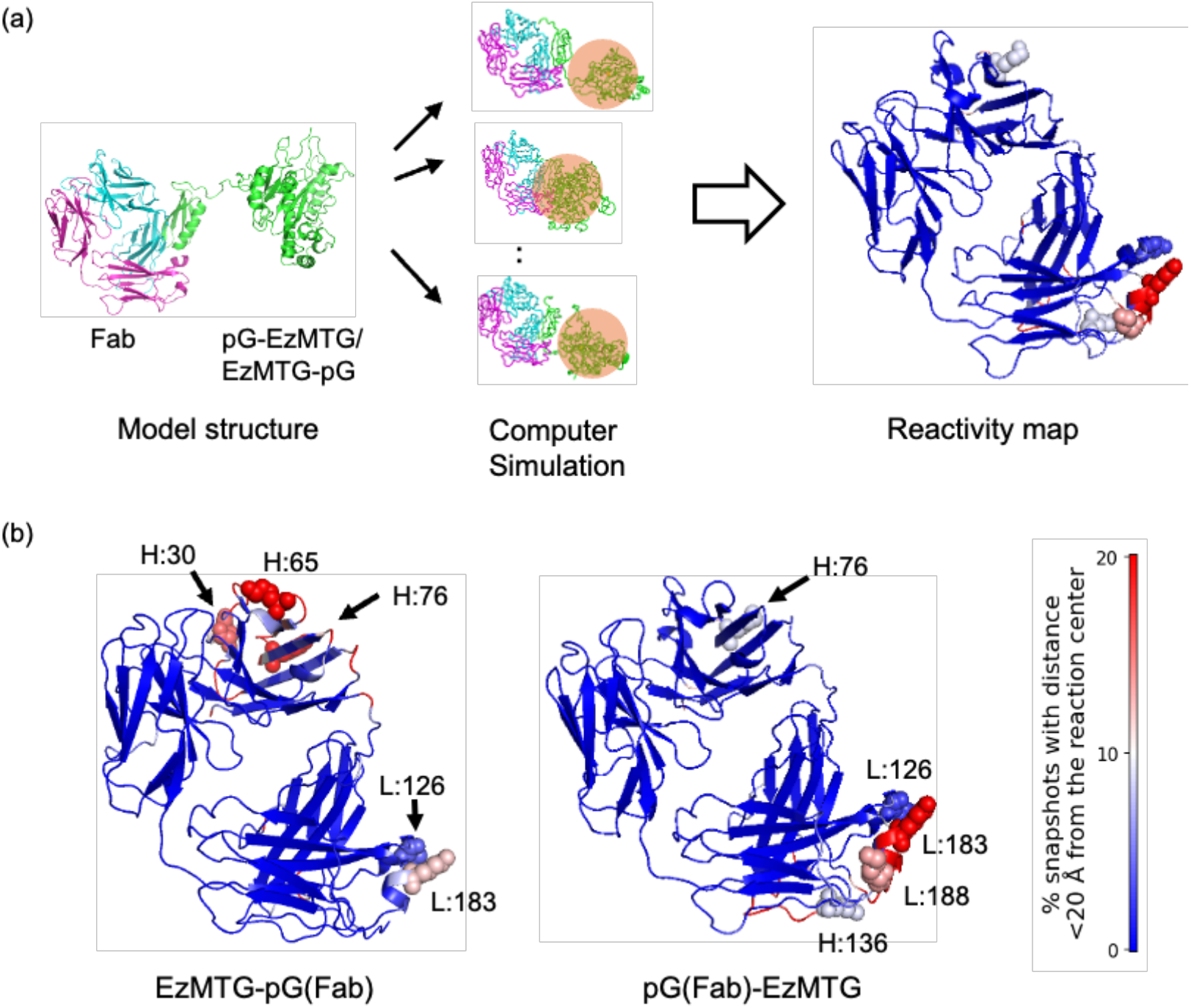
(a) Workflow of the simulation analysis. Representative snapshots of the pG-fused EzMTG and Fab complexes were analyzed to identify residues located within 20 Å of the catalytic center, and encounter frequencies were calculated. (b) Representative structures showing Lys residues frequently located near the catalytic center, with color gradient indicating encounter frequency.

**Fig. S6.**
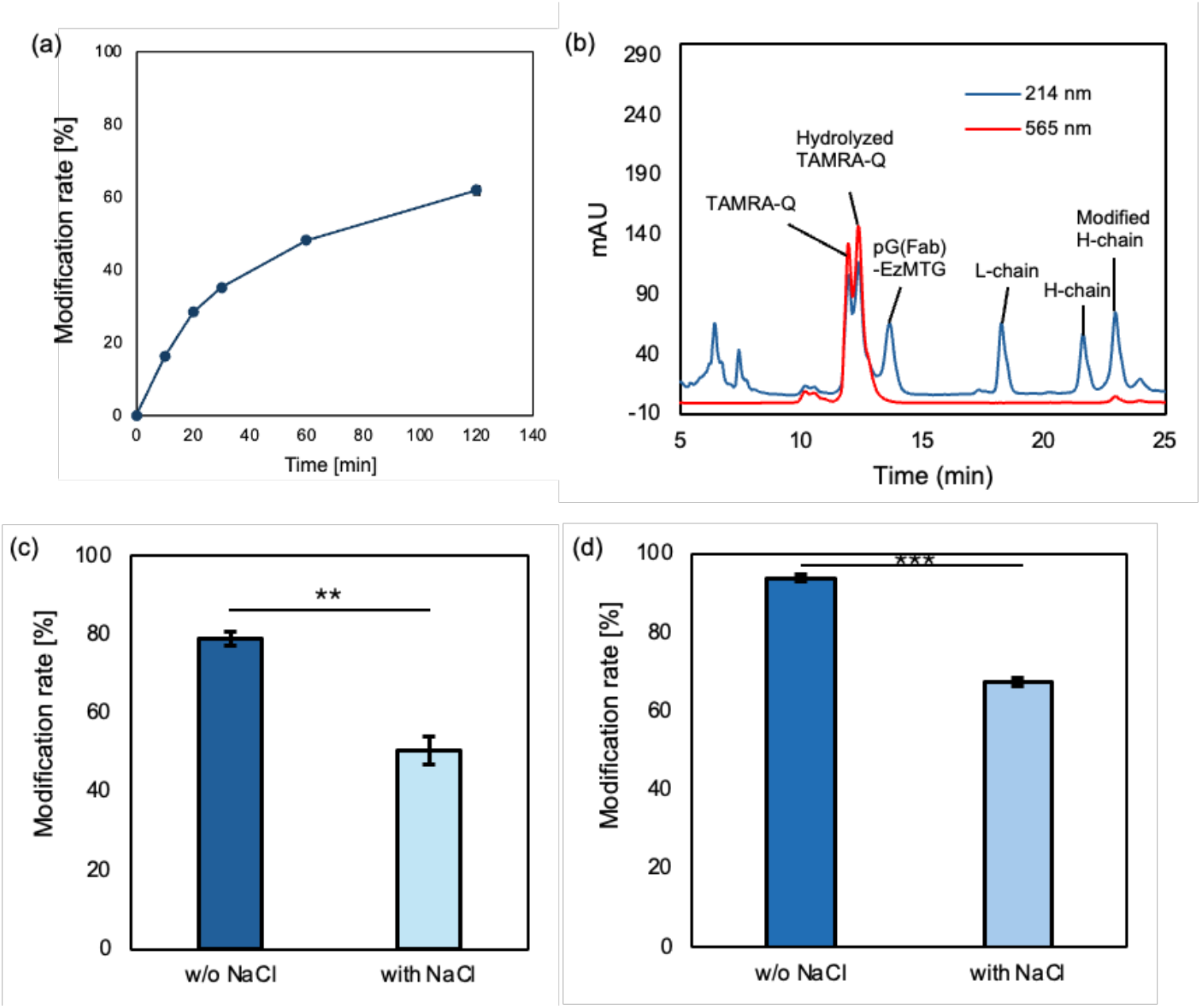
(a) Time course of modification of Lys225 in trastuzumab (1.3 μM) with TAMRA-Q (100 μM) by pG(Fab)-EzMTG (2.6 μM) in 40 mM Tris-HCl (pH 8.0) at 37 °C. *n* = 3; mean ± SE. (b) RP-HPLC chromatograms of reaction solutions with pG(Fab)-EzMTG. Effect of NaCl on trastuzumab modification by (c) pG(Fab)-HA-EzMTG and (d) pG(Fab)-HA(Y62D/P63A)-EzMTG. *n* = 3; mean ± SE; \*\**p* < 0.01, \*\*\**p* < 0.001. The reaction was conducted using trastuzumab (1.3 μM), pG(Fab)-HA-EzMTG (2.6 μM), and TAMRA-Q (100 μM) in 40 mM Tris-HCl (without or with 0.15 M NaCl, pH 8.0) at 37 °C for 120 min.

**Fig. S7.**
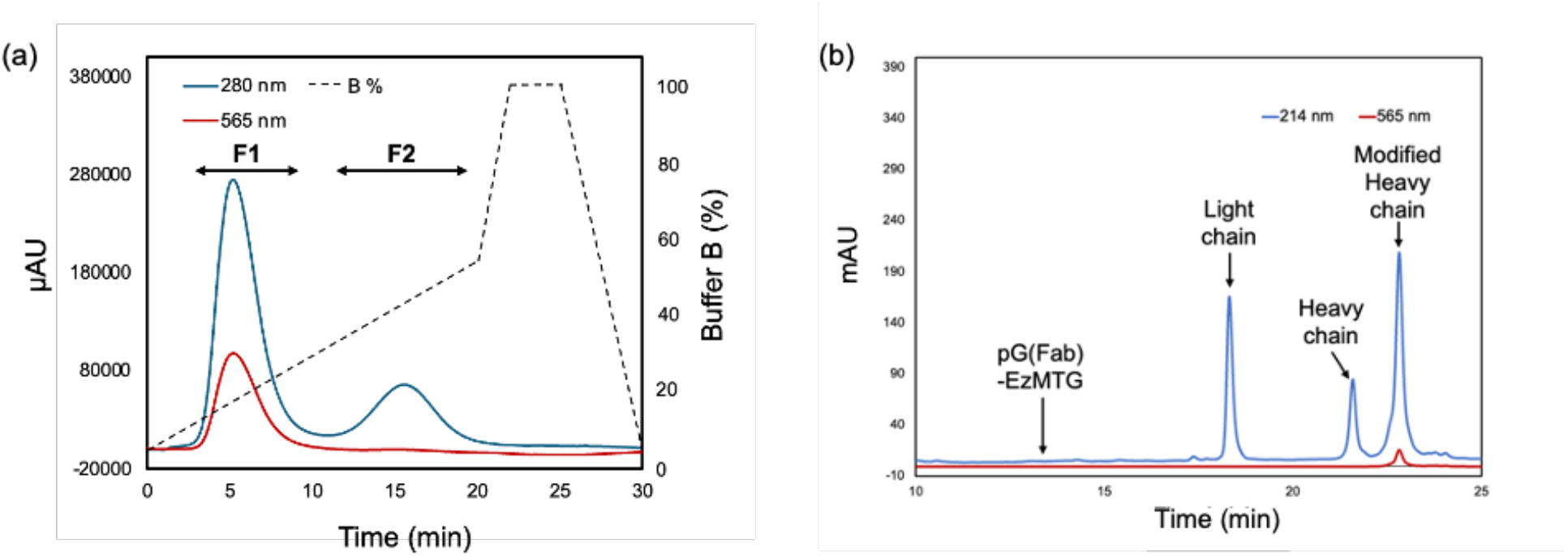
(a) Cation-exchange chromatogram for the purification of TAMRA-Q-modified trastuzumab. Fraction 1 (**F1**) contained TAMRA-Q-modified and unmodified trastuzumab, whereas fraction 2 (**F2**) contained mainly pG(Fab)-HA(Y62D/P63A)-EzMTG. (b) RP-HPLC chromatogram of **F1**, used as purified TAMRA-Q-modified trastuzumab for evaluation of antigen-binding properties.

**Fig. S8.**
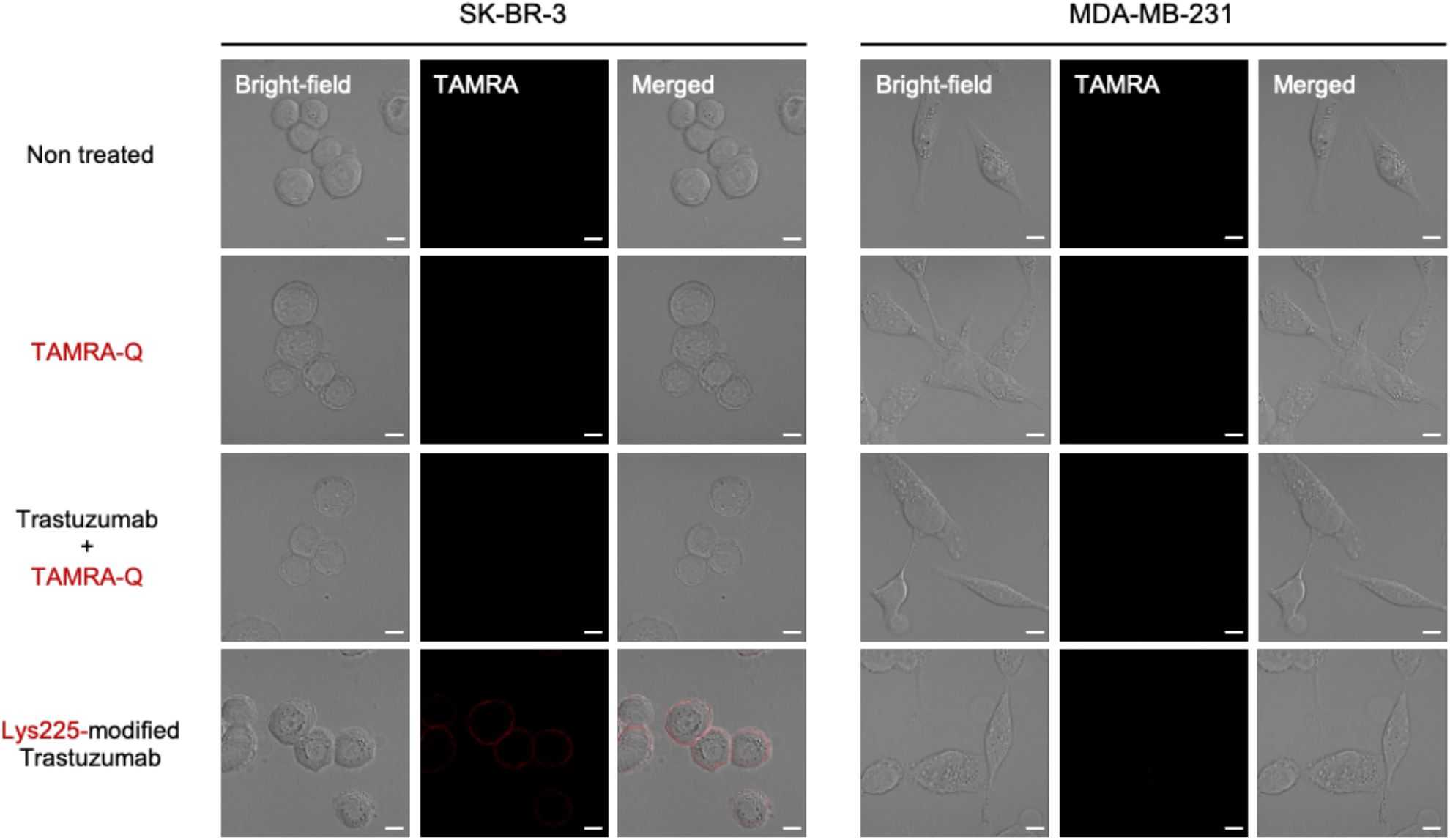
Binding of TAMRA-Q-modified trastuzumab to SK-BR-3 and MDA-MB-231 cell lines. Representative fluorescence images obtained by CLSM. SK-BR-3 and MDA-MB-231 cells were incubated with each sample at 37 °C for 1 h. Bars: 10 μm.

**Fig. S9.**
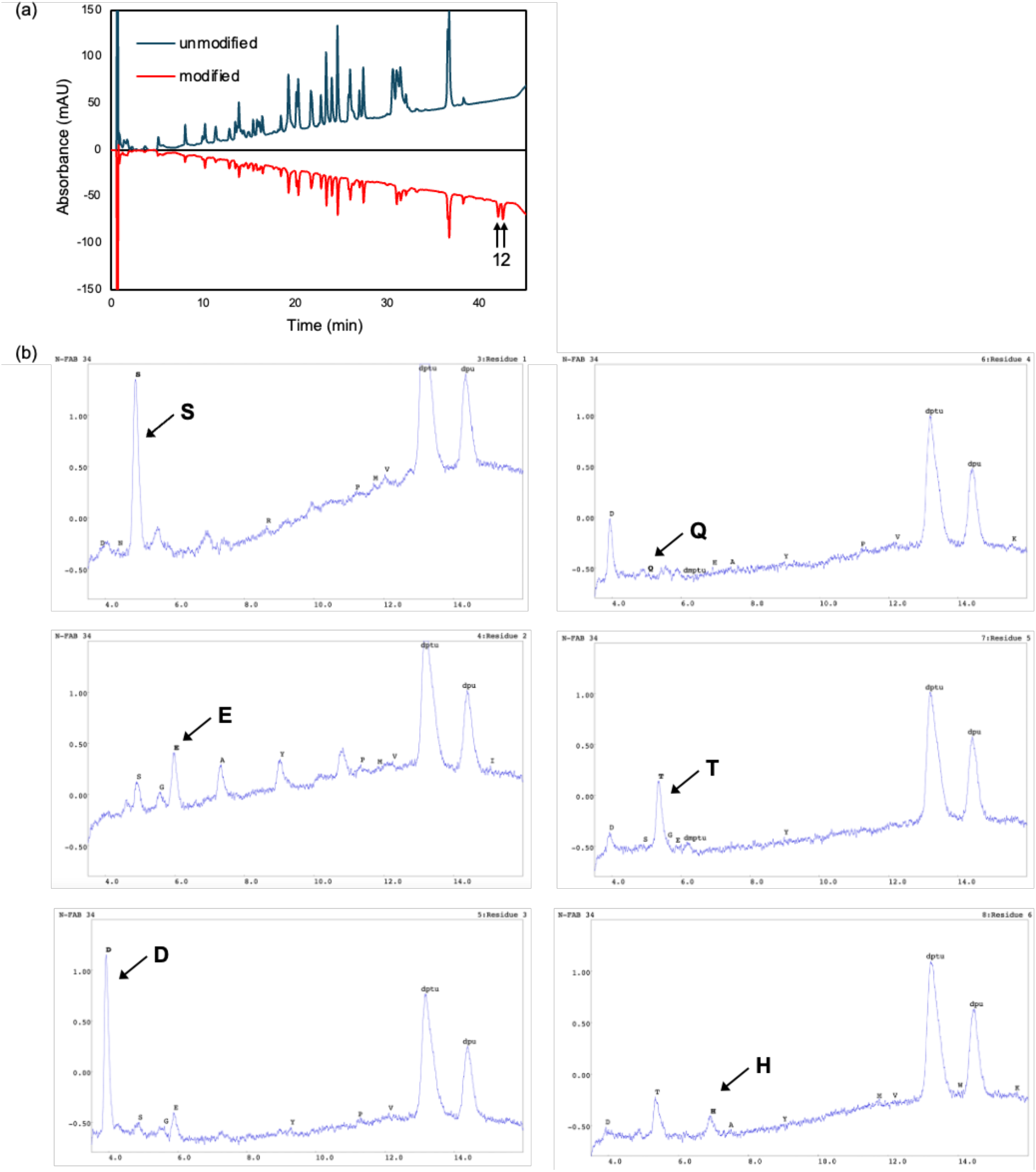
Identification of the modification site using MMAE-Q peptide. (a) RP-HPLC chromatogram of the antibody modified with MMAE-Q peptide by pG(Fab)-HA(Y62D/P63A)-EzMTG. (b) HPLC chromatograms of each residue obtained from *N*-terminal sequence analysis of modified antibody fraction No. 1 in the chromatogram in (a). The same analysis was performed for fraction No. 2, yielding an identical sequence. The results for the second and fourth residues were ambiguous, likely because the second residue corresponds to an alkylated Cys and the fourth residue to a chemically modified Lys.

**Fig. S10.**
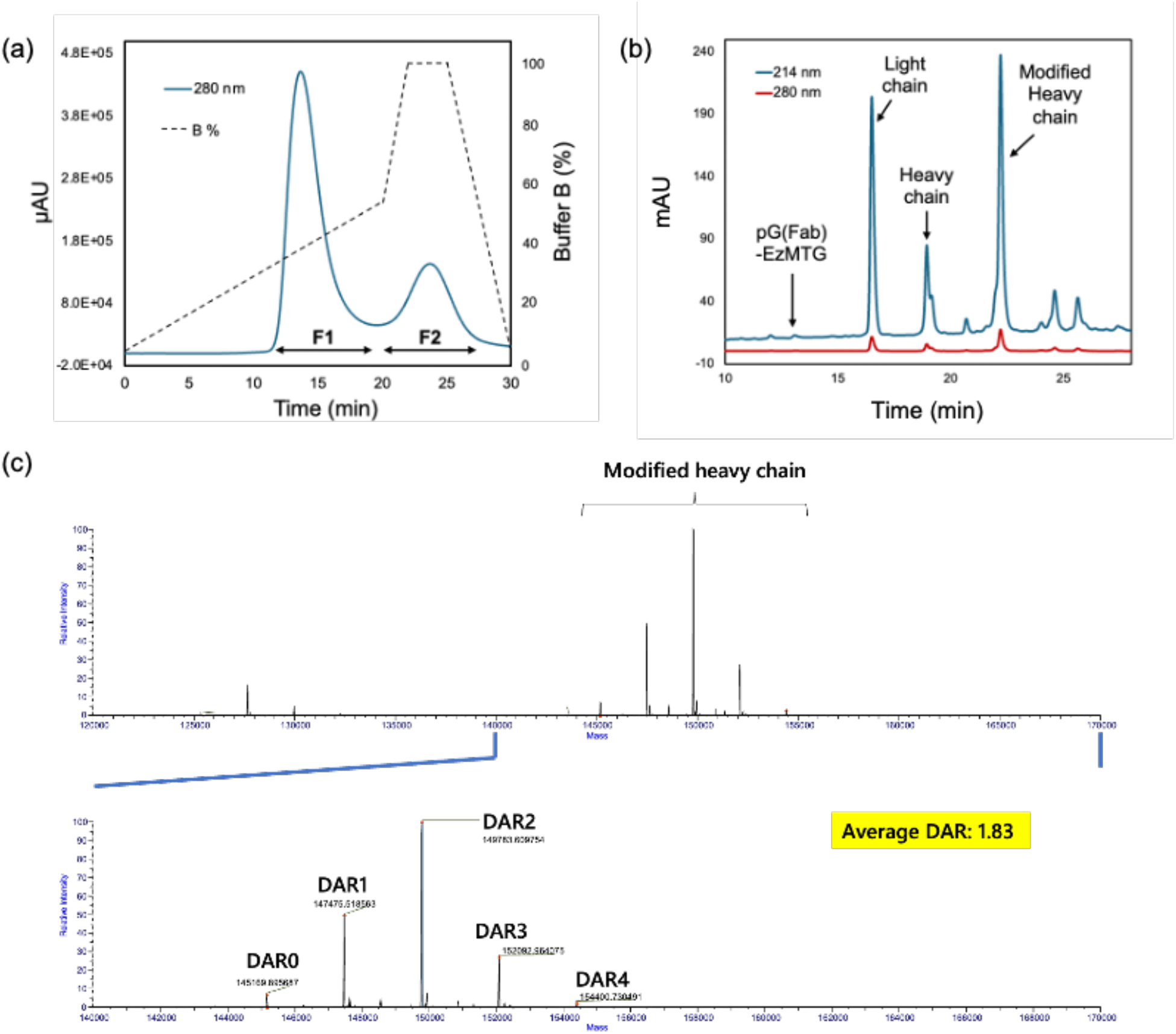
(a) Cation-exchange chromatogram for the purification of MMAE-Q-modified trastuzumab. Fraction 1 (**F1**) contained MMAE-Q-modified and unmodified trastuzumab, whereas fraction 2 (**F2**) contained mainly pG(Fab)-HA(Y62D/P63A)-EzMTG. (b) RP-HPLC chromatogram of **F1**, used as purified MMAE-Q-modified trastuzumab for evaluation of cytotoxicity. (c) LC–MS chromatograms obtained from DAR analysis of the prepared ADC.

---

**Table S1.**
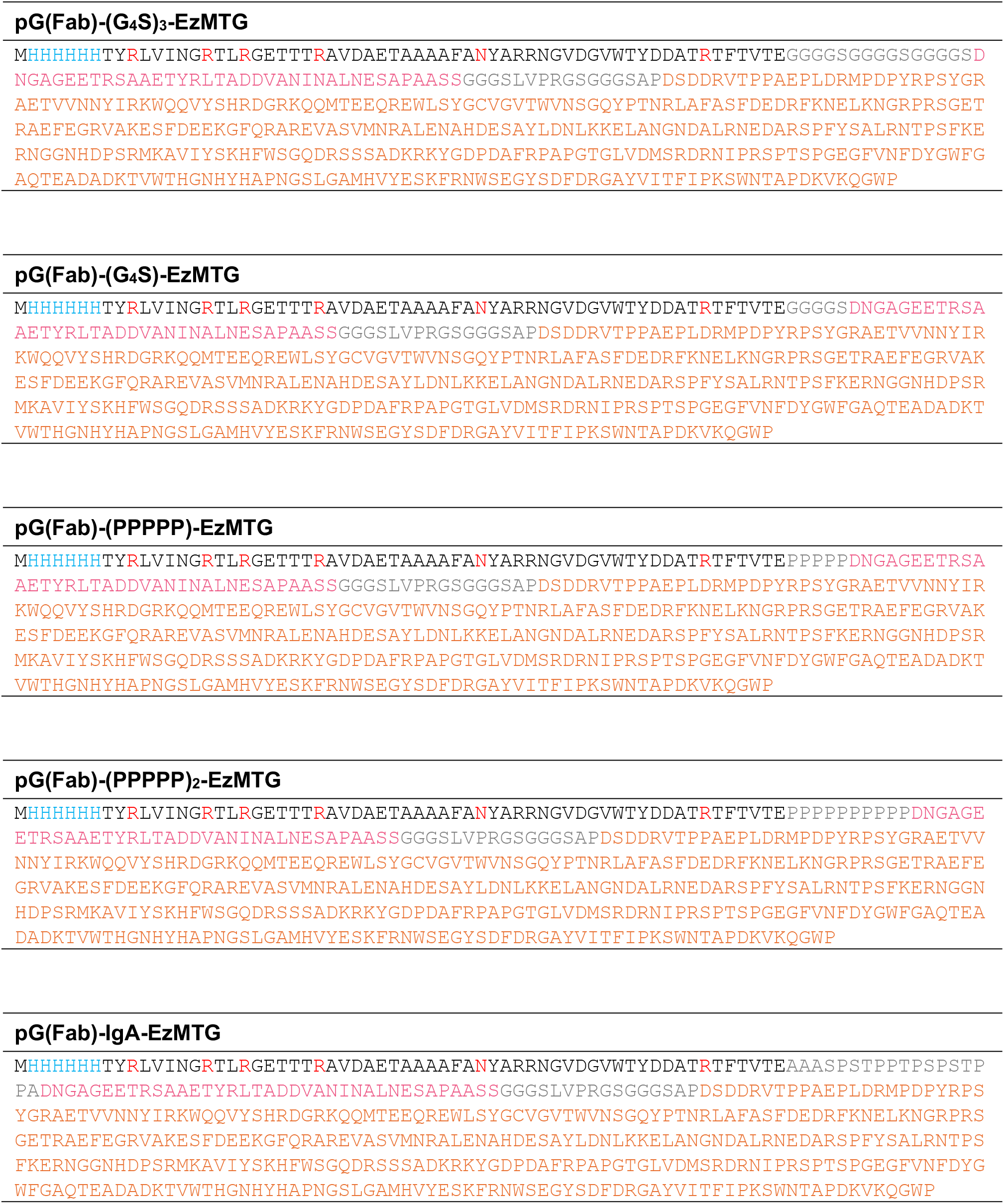
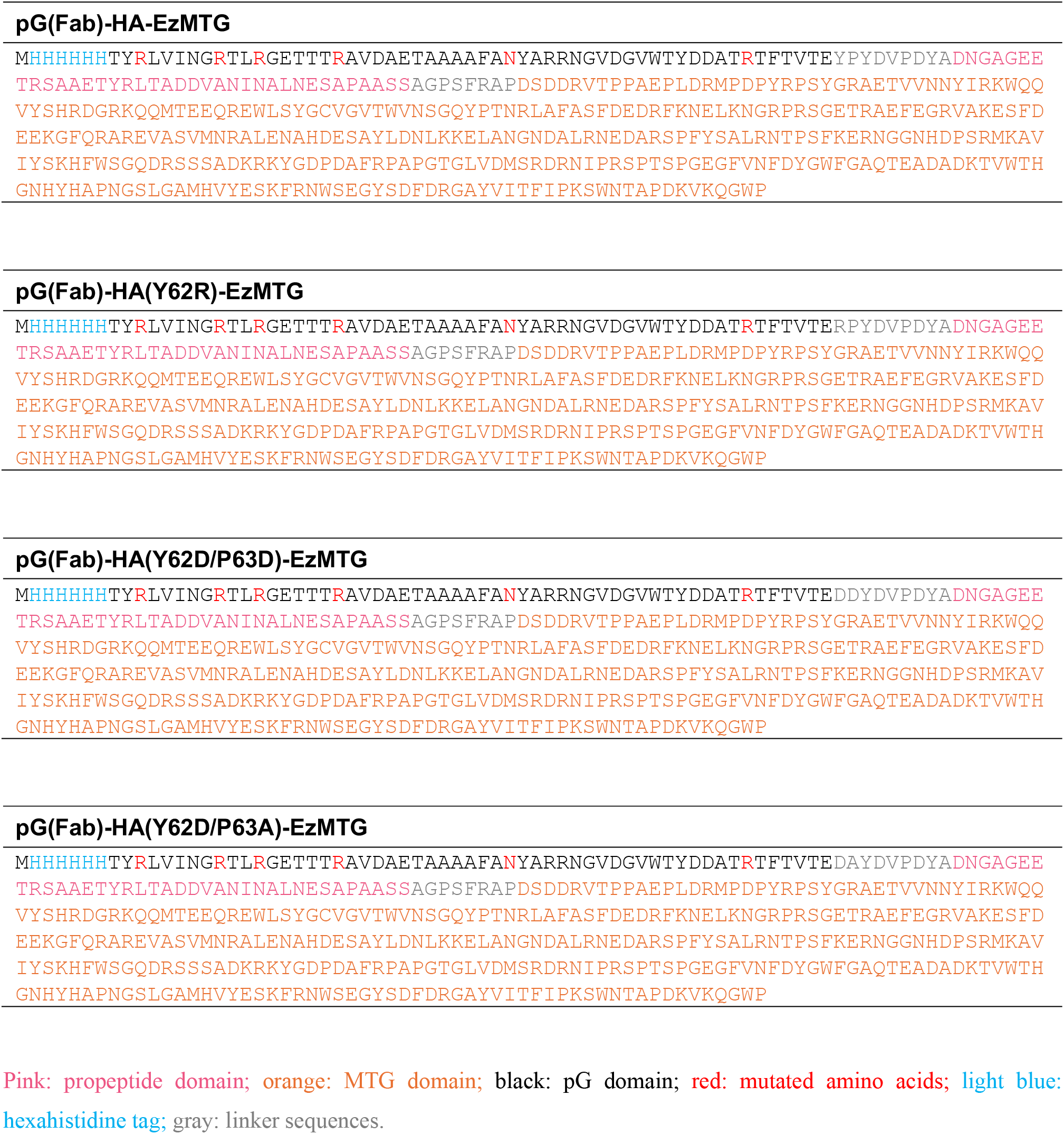
The amino acid sequences of pG-fused EzMTG mutants.

**Table S2.** Lysine residues predicted by simulation to have high reactivity with each pG-fused EzMTG. EzMTG-pG(Fab) pG(Fab)-EzMTG.

| EzMTG-pG(Fab) |  |  | pG(Fab)-EzMTG |  |
| --- | --- | --- | --- | --- |
|  | chain | sequence | chain | sequence |
| 1 | Heavy chain | TSK<sub>76</sub>NT | Light chain | LSK<sub>183</sub>AD |
| 2 | Heavy chain | SVK<sub>65</sub>GR | Light chain | YEK<sub>188</sub>HK |
| 3 | Heavy chain | NIK<sub>30</sub>DT | Heavy chain | SSK<sub>136</sub>ST |

**Table S3.** Characteristics of the lysine residues in the heavy chain of trastuzumab calculated byMolecular Operating Environment (MOE), in descending order of the positive residue patch area (PRPA) value.

| Position | ASA (Å<sup>2</sup>)<sup>1</sup> | Exp (%)<sup>2</sup> | PRPA (Å<sup>2</sup>)<sup>3</sup> |
| --- | --- | --- | --- |
| 65 | 148.5 | 63.06 | 50.4 |
| 225 | 123.4 | 52.41 | 49.1 |
| 249 | 138.7 | 58.91 | 47.8 |
| 363 | 83.4 | 35.4 | 43.2 |
| 208 | 109.7 | 46.6 | 43 |
| 124 | 119.2 | 50.63 | 42.6 |
| 43 | 143.9 | 61.1 | 41.6 |
| 395 | 80.6 | 34.24 | 41.6 |
| 136 | 58.4 | 24.81 | 40.4 |
| 277 | 145.4 | 61.74 | 40.4 |
| 213 | 153.6 | 65.23 | 38.4 |
| 329 | 200.3 | 85.06 | 37.7 |
| 216 | 55.7 | 23.66 | 36 |
| 76 | 152.1 | 64.58 | 35 |
| 325 | 58.2 | 24.74 | 33.8 |
| 291 | 138.1 | 58.66 | 33.4 |
| 337 | 71.2 | 30.25 | 33 |
| 30 | 77.9 | 33.1 | 31.8 |
| 221 | 97.1 | 41.22 | 30 |
| 217 | 94.1 | 39.95 | 28.7 |
| 343 | 126.8 | 53.87 | 28.1 |
| 251 | 42.4 | 18.02 | 27.6 |
| 373 | 21.8 | 9.26 | 22 |
| 341 | 16.6 | 7.07 | 11.1 |
| 293 | 85.7 | 36.41 | 10 |
| 150 | 10.9 | 4.65 | 0 |
| 320 | 78.5 | 33.32 | 0 |
| 323 | 55.8 | 23.7 | 0 |
| 412 | 0.5 | 0.21 | 0 |
| 417 | 44.9 | 19.08 | 0 |
| 442 | 51.6 | 21.9 | 0 |

<sup>1</sup>ASA: accessible surface area of the residue.
<sup>2</sup>Exp: exposure of the residue compared with the ideal surface as determined from Gly\_X\_Gly triplets.
<sup>3</sup>PRPA: positive residue patch area. When the surrounding region possesses an appreciable positive charge, and the residue contributes to positive charges exposed on the surface, the value is increased.

**Table S4.**
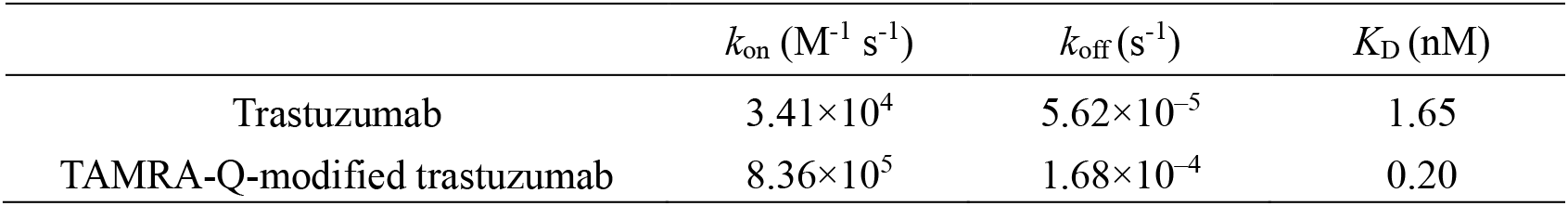
Kinetic parameters for the binding of trastuzumab and TAMRA-Q-modified trastuzumab to HER2 determined using SPR.

| | $k_{\text{on}}$ ($\text{M}^{-1} \text{s}^{-1}$) | $k_{\text{off}}$ ($\text{s}^{-1}$) | $K_{\text{D}}$ (nM) |
| --- | --- | --- | --- |
| Trastuzumab | $3.41 \times 10^4$ | $5.62 \times 10^{-5}$ | 1.65 |
| TAMRA-Q-modified trastuzumab | $8.36 \times 10^5$ | $1.68 \times 10^{-4}$ | 0.20 |

## Notes

### Competing Interest Statement

The authors have declared no competing interest.

